# Nanoparticle Cellular Internalization is Not Required for RNA Delivery to Mature Plant Leaves

**DOI:** 10.1101/2021.03.17.435888

**Authors:** Huan Zhang, Natalie S. Goh, Jeffrey Wang, Gozde S. Demirer, Salwan Butrus, So-Jung Park, Markita P. Landry

## Abstract

Rapidly growing interest in nanoparticle-mediated delivery of DNA and RNA to plants requires a better understanding of how nanoparticles and their cargoes translocate in plant tissues and into plant cells. However, little is known about how the size and shape of nanoparticles influences transport in plants and use of their cargoes, limiting development and deployment of nanotechnology in plant systems. Here, we employ non-biolistically delivered DNA-modified gold nanoparticles (AuNP) spanning various sizes (5 – 20 nm) and shapes (spheres and rods) to systematically investigate their transport following infiltration into *Nicotiana benthamiana (Nb)* leaves. Generally, smaller AuNPs demonstrate more rapid, higher, and longer-lasting levels of association with plant cell walls compared to larger AuNPs. We observe internalization of rod-shaped but not spherical AuNPs into plant cells, yet surprisingly, 10 nm spherical AuNP functionalized with small-interfering RNA (siRNA) are most efficient at siRNA delivery and inducing gene silencing in mature plant leaves. These results indicate the importance of nanoparticle size in efficient biomolecule delivery, and, counterintuitively, demonstrate that efficient cargo delivery is possible and potentially optimal in the absence of nanoparticle cellular internalization. Our results highlight nanoparticle features of importance for transport within plant tissues, providing a mechanistic overview of how nanoparticles can be designed to achieve efficacious bio-cargo delivery for future developments in plant nanobiotechnology.

## Introduction

The growth of nanobiotechnology, whereby nanomaterials are designed for use in biological systems, has added new dimensionality to pharmaceutical and drug delivery development and married the fields of chemistry, biomedical engineering, and material science. Their small size, tunable physicochemical, unique optical properties, and high surface-to-volume ratio render nanoparticles (NPs) versatile scaffolds to be functionalized as carriers or probes for therapeutic and diagnostic purposes^1–3^. In recent years, the use of nanomaterials in plant science has greatly enabled advances in agriculture. These developments include crop management improvement, plant-pathogen protection, nutrient and pesticide delivery, monitoring of plant and soil health, and creation of crop varieties with desirable traits such as high-yield and stress resistance^4–6^. In plant genetic engineering, nanomaterials have also been used as vehicles for the delivery of plasmid DNA^7–9^, siRNA^10–12^, and proteins^13^ to whole plants.

Nanoparticle (NP)-mediated biomolecule delivery technologies for plants have leveraged the material properties of nanoparticles to overcome the unique barriers of plant cells. Examples of these materials include mesoporous silica nanoparticle (MSNs)^7^, single walled carbon nanotubes (SWNTs)^8,9,11,14^, DNA nanostructures^12^, and layered double hydroxide nanosheets^10,15^. These works exploit the small size and high tensile strength of NPs to bypass biological barriers, the diverse conjugation chemistries available for cargo conjugation, and the high degree of control over morphology and surface functionalization to provide better penetration through plant tissues. While NP-mediated biomolecule delivery in plants has been demonstrated with nanoparticles of various sizes and surface modifications, little is known about how these design variables affect translocation, cellular uptake, and ultimately bio-cargo utilization in plants. Recent studies allude that NP size, surface charge, and stiffness are important factors for NP transport across the cell walls in plants^12,16^. Comprehensive studies in mammalian systems^17–20^ and simulations of NP-cell membrane interactions^21–24^ have underscored the importance of NP size and shape for bio-delivery – yet, these studies are usually not applicable to walled plant cells. In particular, studies in mammalian systems have revealed how size and shape can affect NP interaction with cells and thus influence the uptake pathway and internalization efficiency, guiding design strategies for nanoparticle-based biomedical applications^25^. However, analogous studies in plant systems are uncommon, limiting the development of plant nanobiotechnologies and the assessment of their intended and unintended impacts on plants, agricultural systems, and the environment.

The plant cell wall is composed of a complex network of biopolymers that gives rise to a semipermeable matrix^26^ and is a unique barrier to consider for the uptake of NPs into plant cells^27^. While the permeability of the plant cell wall is dynamic, studies commonly suggest that the upper limit of the pore diameters range from 5 to 20 nm^28,29^. An abundance of studies in mammalian systems have demonstrated that NP interactions with cells is morphology-dependent, and similar dependencies may also dictate NP interactions with plants. There is therefore a cogent need to understand NP-plant interactions to inform how NPs can be designed for use with minimal environmental disruption and maximal efficacy. The process of NP uptake, translocation, and accumulation in plants can be broadly split into three tiers: (i) macroscale – quantifying translocation and accumulation in plant organs, (ii) microscale – studying NP transport through and interactions with plant tissues and vasculature, and (iii) molecular – revealing the manner of NP association on a cellular or sub-cellular level^30^. Of note, most studies have been performed on the macroscale^31,32^, though some studies have begun to explore the effect of NP properties on uptake on the microscale^33,34^. While these studies are valuable to understand how NPs and their surface chemistries might impact long-term accumulation and translocation throughout a plant, NP features enabling molecular-scale translocation of NPs within plant tissues and into plant cells, and subsequent bio-cargo delivery, remain to be determined.

In this study, we design a model nanoparticle system based on DNA-functionalized gold nanoparticles (AuNP) of varied sizes (5, 10, 15, and 20 nm) and shapes (spheres and rods, denoted as AuNS and AuNR, respectively). While AuNP are broadly used for biolistic delivery of biological cargoes in plants, their non-biolistic delivery remains unexplored. Herein, we track individual AuNP and investigate how nanoparticle size and morphology affect nanoparticle interaction with and uptake into plant cells. We utilize confocal microscopy and transmission electron microscopy (TEM) to track AuNP movement and directly visualize NP interactions with plant cells. Our results demonstrate a NP morphology-dependent association and internalization into plant cells in a time-dependent manner. Specifically, colocalization analysis of fluorescently tagged AuNP reveals that smaller AuNSs reach peak association with plant cell walls earlier and at higher levels than larger spherical AuNS. Interestingly, TEM results indicate cellular internalization of AuNP may be shape-dependent; AuNR are identified inside cells but not AuNS of any size. Inhibitor assays suggest the uptake mechanism of the AuNP to be morphology-dependent, whereby AuNR internalize into plant cells through endocytosis, and further support the finding that AuNS do not enter cells. These results suggest that endocytosis is a likely mechanism of AuNR internalization, and support TEM-based observations that AuNS do not internalize into plant cells. Finally, we demonstrate that both AuNS and AuNR can carry siRNA to silence GFP in transgenic GFP *Nicotiana benthamiana* (*Nb*) plants, despite a lack of AuNS cellular internalization. We find that non-internalizing 10 nm AuNS can achieve 99% silencing efficiency on the mRNA level and show the highest levels of silencing amongst all AuNPs. Taken together, our results highlight the importance of nanoparticle morphology in transport within plant tissues and suggest that efficient cellular siRNA delivery can be achieved even without carrier internalization into cells.

## Main

In this study, we uncover the relationship between nanoparticle morphology and degree of association with intact cells of mature plants for downstream applications in siRNA delivery. Gold nanoparticles possess several desirable characteristics that favor them as a model nanoparticle system, including their ease of synthesis over various nanoscale dimensions, well-validated and diverse functionalization chemistries, high contrast under electron microscopy for tracking, low morphological polydispersity, and high biocompatibility^35–37^. Here, we designed a library of DNA-functionalized gold nanoparticles (DNA-AuNPs) of varied morphologies, evaluated the leaf tissue transport of fluorophore-tagged DNA-AuNPs to plant cells over time, established a size and shape-dependent phenomenon for DNA-AuNP transport to and entry through plant cell walls, and demonstrated that bio-cargo delivery into plant cells can be independent of nanoparticle cellular internalization.

### Preparation and characterization of DNA-AuNP

Five gold nanoparticles of various morphologies were used in this study – 5 nm, 10 nm, 15 nm and 20 nm gold nanospheres (AuNS), and a 13 by 68 nm gold nanorod (AuNR) of aspect ratio ∼5.2 (**Fig. 1a**). To obtain particles that are colloidally-stable in plants, AuNP were functionalized with single-stranded DNA sequences that were thiol-modified on the 5’ end, followed by a 10-nucleotide poly-adenosine sequence to enhance 5’ association with the gold nanoparticle surface and, when necessary, increase the availability of the 3’ end of the DNA strand for cargo (fluorophore or siRNA) hybridization^38,39^ (**Table S1**). DNA functionalization was accomplished through a pH-assisted method^40^, which functionalized AuNP with thiol-modified DNA at low pH and low salt concentrations, providing a quick and efficient method of nucleic acid (NA) functionalization in place of the more widely used “salt-aging” technique (functionalization details in **Table S2**). Citrate-stabilized AuNP and DNA-AuNP were characterized with UV-Vis spectroscopy, TEM imaging, and dynamic light scattering (DLS). AuNS have a size-dependent surface plasmon resonance (SPR) peak at approximately 520 nm and AuNR have characteristic transverse and longitudinal peaks at 520 nm and 960 nm (**Fig. 1b** and **Fig. S1**). Successful functionalization of AuNP was confirmed using DLS measurements, appropriate for spherical AuNS and DNA-AuNS, which indicate DNA functionalization results in a 9 - 18 nm increase in NP hydrodynamic size (**Table S3** and **Fig. S2** and **S3**, AuNR were not analyzed due to inherent limitations of DLS with non-spherical particles). Further confirmation included UV-Vis spectroscopy, whereby DNA-AuNS experience a small redshift in SPR peak compared to their citrate-stabilized counterparts^41^ (**Fig. S1**). TEM images of AuNP (**Fig. 1c**) and DNA-AuNP (**Fig. S4**) demonstrate the high homogeneity of AuNP before and after DNA functionalization. We optimized and quantified the density of DNA on each of the five DNA-AuNPs with UV-Vis spectroscopy post-KCN treatment^42^. The number of DNA molecules per AuNP are shown in Table S4 with AuNPs possessing higher surface areas loading more DNA, as expected. DNA-AuNP were next tagged with a Cy3 fluorophore and abaxially-infiltrated into mature leaves of *Nicotiana benthamiana* (*Nb*) plants using a needleless syringe (**Fig. 1d**).

**Fig. 1.**
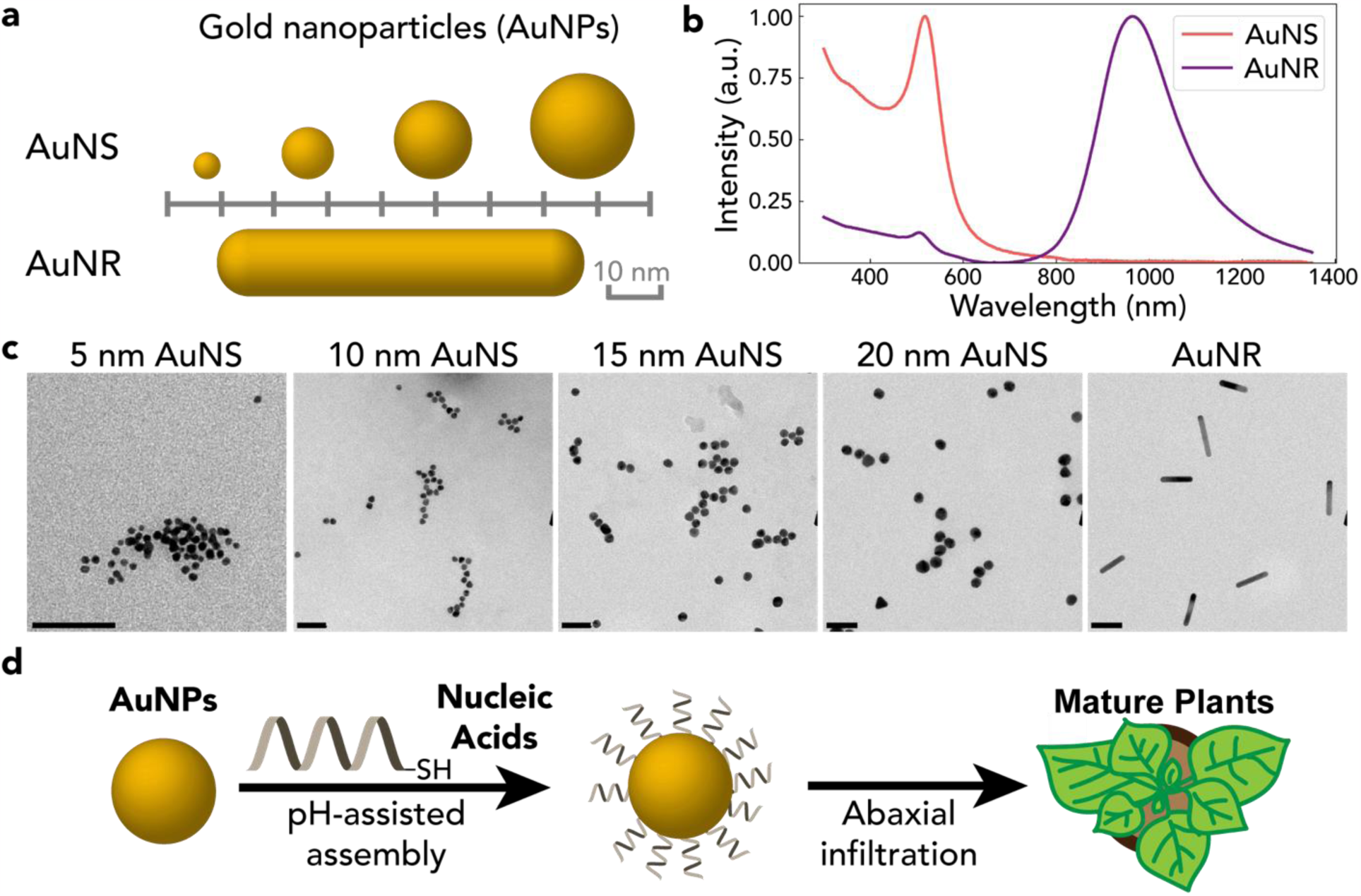
Gold nanoparticle (AuNP) and DNA-AuNP preparation and characterization. **a** Illustration of the five AuNPs used in this study: 5 nm, 10 nm, 15 nm, and 20 nm gold nanospheres (AuNS), and 13 by 70 nm gold nanorods (AuNR). **b** Representative visible-near IR spectra of 10 nm AuNS and AuNR demonstrate characteristic AuNS and AuNR absorbance peaks. **c** TEM characterization of AuNP indicates high degrees of monodispersity and structural stability due to the uniform shape of AuNP as seen in these representative images. Scale bar: 50 nm. **d** AuNP are functionalized with thiol-modified nucleic acids via a low-pH assisted method, followed by infiltration into mature plant leaves for downstream studies.

### Morphology and time dependence of AuNP transport through leaf tissue and association with plant cells

Following nanoparticle characterization, we tested the morphology- and size-dependent ability of AuNPs to transit within the leaf interstitial space and associate with plant cell walls. To track DNA-AuNP in plant tissue, we fluorescently-tagged the AuNP by hybridizing a Cy3-modified DNA strand containing a sequence complementary to the thiol-modified DNA used to functionalize AuNP (sequence in **Table S1**). Cy3-labelled DNA-AuNP (Cy3-DNA-AuNP) with a normalized Cy3 concentration of 400 nM were infiltrated into the leaves of a transgenic *Nb* plant that constitutively expresses mGFP5, incubated for a set amount of time post-infiltration, and the infiltrated leaf segments were mounted and imaged with confocal microscopy. Signal from Cy3 and GFP channels was collected, whereby Cy3 signal enables Cy3-DNA-AuNP localization and GFP from the transgenic *Nb* line provides a fluorescent marker originating from the intracellular space of the cell. The colocalization fraction generated by overlaying Cy3 and GFP channels was analyzed to evaluate the relative ability of Cy3-DNA-AuNP to diffuse within the leaf interstitial space and associate with the plant cells (Fig. 2a).

**Fig. 2.**
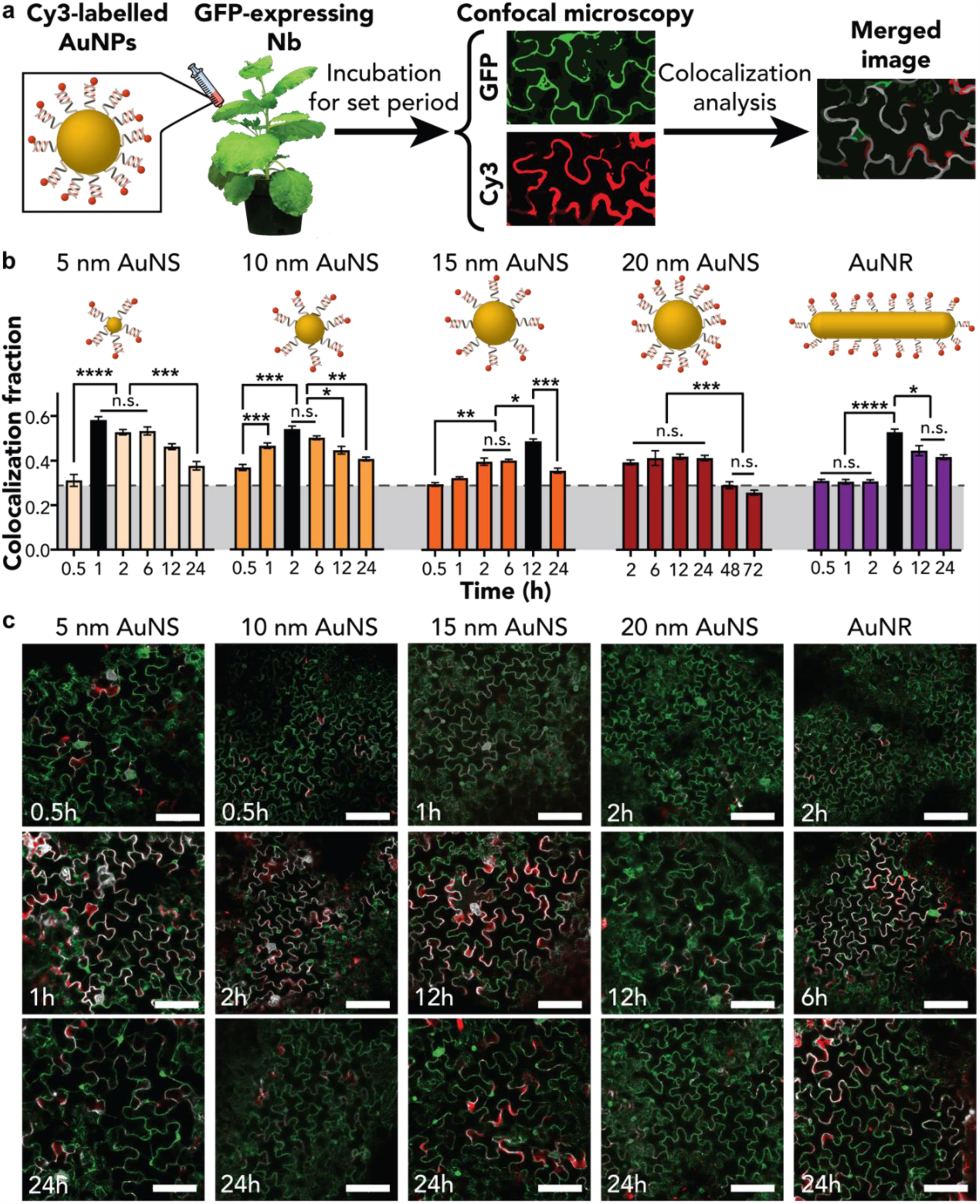
Cy3-tagged DNA-AuNP association with plant cells following infiltration into transgenic mGFP5 *Nicotiana benthamiana* (*Nb*) leaves. **a** Workflow for fluorescent AuNP tracking in leaves via confocal microscopy. Cy3-labelled DNA hybridized with a complementary sequence on DNA-functionalized AuNP, infiltrated into mGFP5 *Nb* leaves, and analyzed under confocal microscopy after a defined incubation period. Constitutively expressed GFP fluorescence serves as the intracellular cytoplasm marker, and the colocalization fraction between the Cy3 and GFP channels serve as a relative measure of AuNP cell wall intercalation and cell internalization. **b** Time-series of colocalization fractions for Cy3-DNA-AuNP demonstrate size and shape dependent AuNP intercalation and by extension, a similar trend for internalization. Colocalization values increase gradually and reach respective maxima at shorter timepoints (1h for 5 nm, 2h for 10 nm, and 12h for 15nm) for smaller AuNS sizes, while larger sized AuNS (20 nm) do not achieve a clear maximum colocalization value at any timepoints assayed, suggesting that AuNS above 20 nm in size may be less amenable to transport within leaf tissue and ultimately internalization. Interestingly, AuNR infiltrated leaves show a sudden increase to peak colocalization at 6h. Grey denotes assay background levels of detection with a free Cy3-DNA control. (5nm AuNS, ****p<0.0001, ***p=0.0008; 10nm AuNS: ***p=0.0003, **p=0.0030, *p=0.0357; 15nm AuNS: **p=0.0056, *p=0.0276, ***p=0.0009; 20nm AuNS: ***p=0.0004; AuNR: ****p<0.0001, *p=0.0115 in one-way ANOVA, n.s.: not significant; error bars indicate s.e.m. n = 3). **c** Representative confocal images of Cy3-DNA-AuNP infiltrated *Nb* leaves with regions of colocalization (white) between intracellular GFP (green) and Cy3 (red) channels. Images corresponding to incubation times included for each of the five AuNPs include before, during, and after the time for co-localization values to reach the maximum level. Scale bar: 100 µm.

Colocalization analysis of Cy3-DNA-AuNP across timepoints ranging from 30 minutes to 24 hours post-infiltration reveal maximum colocalization fractions at varied timepoints that depend on NP core shape and size (**Fig. 2b**, maxima: black bars), with the exception of 20 nm AuNS, for which no maximum was observed. For DNA-AuNS, the time required to reach maximum AuNP colocalization with plant cells occurred gradually and was shorter for smaller nanoparticles – for 5, 10, and 15 nm, the time corresponding to the colocalization maxima were 1, 2, and 12 hours post-infiltration. For 20 nm AuNS, we did not observe a statistically-significant change in the colocalization fractions (no clear maxima) within 24 hours. Furthermore, longer post-infiltration times of 48 and 72 hours did not yield a notable maximum value for the 20 nm AuNS. For DNA-AuNR, the maximum colocalization fraction occurred 6 hours post-infiltration, attained sharply compared to earlier timepoints. Confocal images in **Fig. 2c** provide a visual representation for the colocalization analysis where Cy3-DNA-AuNP signal was collected as the red channel, intracellular GFP as the green channel, and overlapping signal from the two channels represented in white. With the exception of the 20 nm AuNS, all AuNPs experienced an increase in colocalization fractions as a function of time post-infiltration before reaching a maximum, followed by a decrease in colocalization values over longer incubation times (representative images are shown in **Fig. S5 to S9**). We attribute the time-dependent increase in signal to the time required for AuNPs to travel through plant tissue and intercalate between cells. Interestingly, the time-dependent accumulation of AuNS occurs gradually, whereas time-dependent accumulation of AuNR occurs suddenly at 6 hours post-infiltration, suggesting a shape-dependent effect on NP transport in plant tissues. Meanwhile, the decrease in Cy3 signal at longer incubation points is potentially due to the loss of Cy3 fluorescence with photobleaching or quenching by molecular interactions, or apoplastic transport decreasing Cy3 presence within the imaged area^43^. Our data suggest that AuNS core size plays an essential role for DNA-AuNP transport in plant tissues, with times for maximal cellular association ranging drastically from 1 to 12 hours, and further supports prior hypothetical observations that larger nanoparticles take longer to diffuse through plant tissue and associate with plant cells, with an upper 20 nm NP core size limit.

In sum, our colocalization results suggest that 5, 10, and 15 nm DNA-AuNS and AuNR can diffuse within the leaf interstitial space and associate with plant cell walls and membranes in a size and shape-dependent manner. To confirm NP fate on a sub-cellular scale, we next implemented TEM of NP-treated plant leaves.

### Direct visualization of AuNS association with plant leaf cells

To complement our light microscopy results, we exploit the high electron density of gold-based materials to directly visualize AuNP in plants with TEM (**Fig. 3**). DNA-AuNP infiltrated *Nb* leaves were fixed and sectioned 24 h post-infiltration, with representative images at progressively increased magnifications provided for 5, 10, 15, and 20 nm AuNS, with red boxes denoting the area magnified (**Fig. 3a to 3d**). At the greatest magnification (right most column in **Fig. 3**), solid arrows mark AuNS associated with a single cell wall, and unfilled arrows mark AuNS sandwiched between two neighboring cell walls. Characteristic cellular structures like the nucleus and chloroplasts are used as indicators to determine whether AuNS are localized within the extracellular or intracellular spaces.

**Fig. 3.**
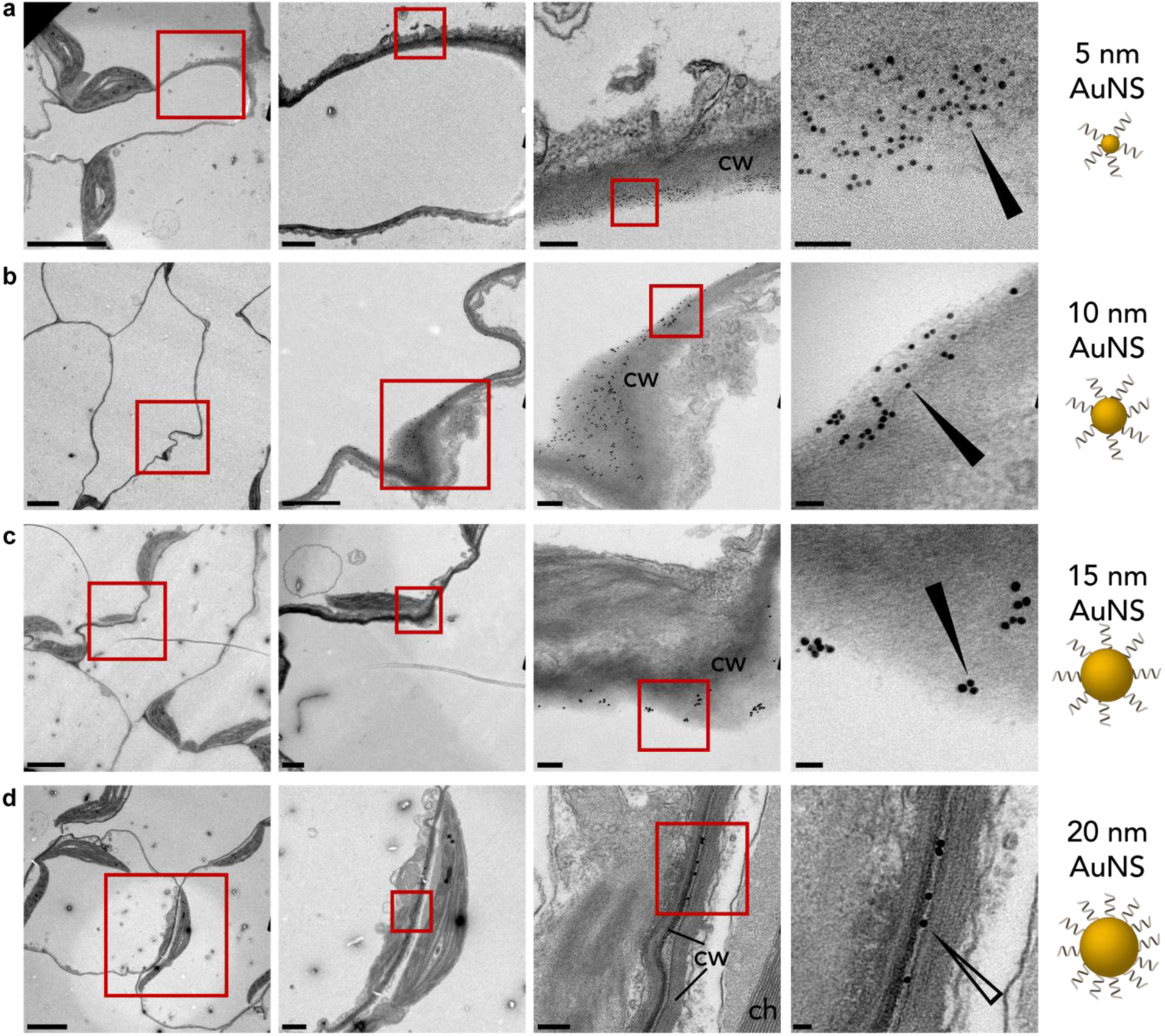
TEM of DNA-functionalized AuNS treated *Nicotiana benthamiana* (*Nb*) leaves. Representative TEM images of *Nb* plants 24h post-infiltration with DNA-functionalized AuNS of **a** 5 nm, **b** 10 nm, **c** 15 nm, and **d** 20 nm diameters. Strong AuNS intercalation between plant cell walls are apparent for AuNS sized 5 nm – 15 nm only. Conversely, 20 nm AuNS were predominantly found between the cell walls of adjacent plant cells. Annotations represent cell wall (cw) and chloroplast (ch). Solid and unfilled arrows indicate nanoparticles associated with a single cell wall or found between cell walls, respectively. Scale bars (from left to right) are 5 µm, 1 µm, 0.2 µm and 50 nm.

Across all leaves infiltrated with different AuNS sizes and concentrations (**Table S4**), a high density of AuNS were identified associated with the plant cell walls, with smaller AuNS associating more than larger AuNS. We hypothesize that the size of the AuNP affects their translocation efficiency within plant tissues, whereby larger AuNPs experience greater difficulty bypassing biological barriers like the cuticle or epidermal layers to access inner leaf layers like the mesophyll. Notably, 5, 10, and 15 nm DNA-AuNS TEM images showed AuNS associated with and intercalating into a single cell wall, whereas the 20 nm AuNS images largely depicted AuNS sandwiched between two cell walls and thus unassociated with a single cell (**Fig. 3d**, more images in **Fig. S10**). Overall, we saw no instances of AuNS within or proximal to the intracellular space of plant cells, suggesting that AuNS internalization into plant cells is minimal, if it occurs at all. The higher degree of plant leaf cell association of smaller AuNP based on TEM agree with their respective higher colocalization values obtained with our confocal microscopy results.

### Direct visualization of AuNR association with and internalization into plant leaf cells

Analogous to the AuNS TEM experiments, *Nb* leaves exposed to DNA-AuNR were prepared for TEM analysis 24 h post-infiltration. In contrast to AuNS of all sizes, we identified several instances of AuNR inside plant cells, with representative images in **Fig. 4a**. The striped arrow pinpoints AuNR found within plant cells, supported by the presence of chloroplasts next to the identified AuNR. In addition, several instances of AuNR piercing into the cell walls were observed (**Fig. 4b** and **Fig. S11**). We further observed that AuNR can orient along a tangent or perpendicular to the cell wall (**Fig. 4b**, **Fig. S11**). Previous theoretical and experimental studies of NP internalization in mammalian cells indicate that NP shape greatly influences the contact curvature with the lipid membrane and dictates the endocytic pathway and angle of entry^23^. In particular, one-dimensional NPs tend to rotate to orient themselves perpendicular to the cell membrane during internalization into mammalian cells^21^, and the rotation can facilitate the penetration and transport of NPs in mucosal tissue^44^. From our TEM data, we therefore analyzed 324 AuNR in the extracellular space of plant cells with respect to their orientation relative to the cell wall tangent (**Fig. 4c**, **Fig. S10**). In total, 41.7%, 21.9%, and 36.4% of AuNR oriented between 0° - 30°, 30° - 60°, and 60° - 90° with the cell wall respectively (details in **Table S4**). We posit that the AuNR demonstrate a stronger preference for initially orienting parallel to the cell wall, where 22.5% of AuNR formed an acute angle between 0°- 10° with the cell wall. In addition, compared to the 11.1% proportion expected randomly, 14.2% of AuNR were oriented between 80° - 90° relative to the plane of the cell wall. Several AuNR that were not oriented in parallel with the cell wall also demonstrated some travel past the cell wall interface (**Fig.4b** and **Fig. S11**). Considering the rigid and porous structure of the cell wall, our data suggest that AuNR experience re-orientations to bypass the cell wall and membrane for plant cell internalization, similar to the phenomenon previously identified in mammalian systems suggesting that asymmetric nanoparticles are entropically favored for membrane interactions^21,23,44^.

**Fig. 4.**
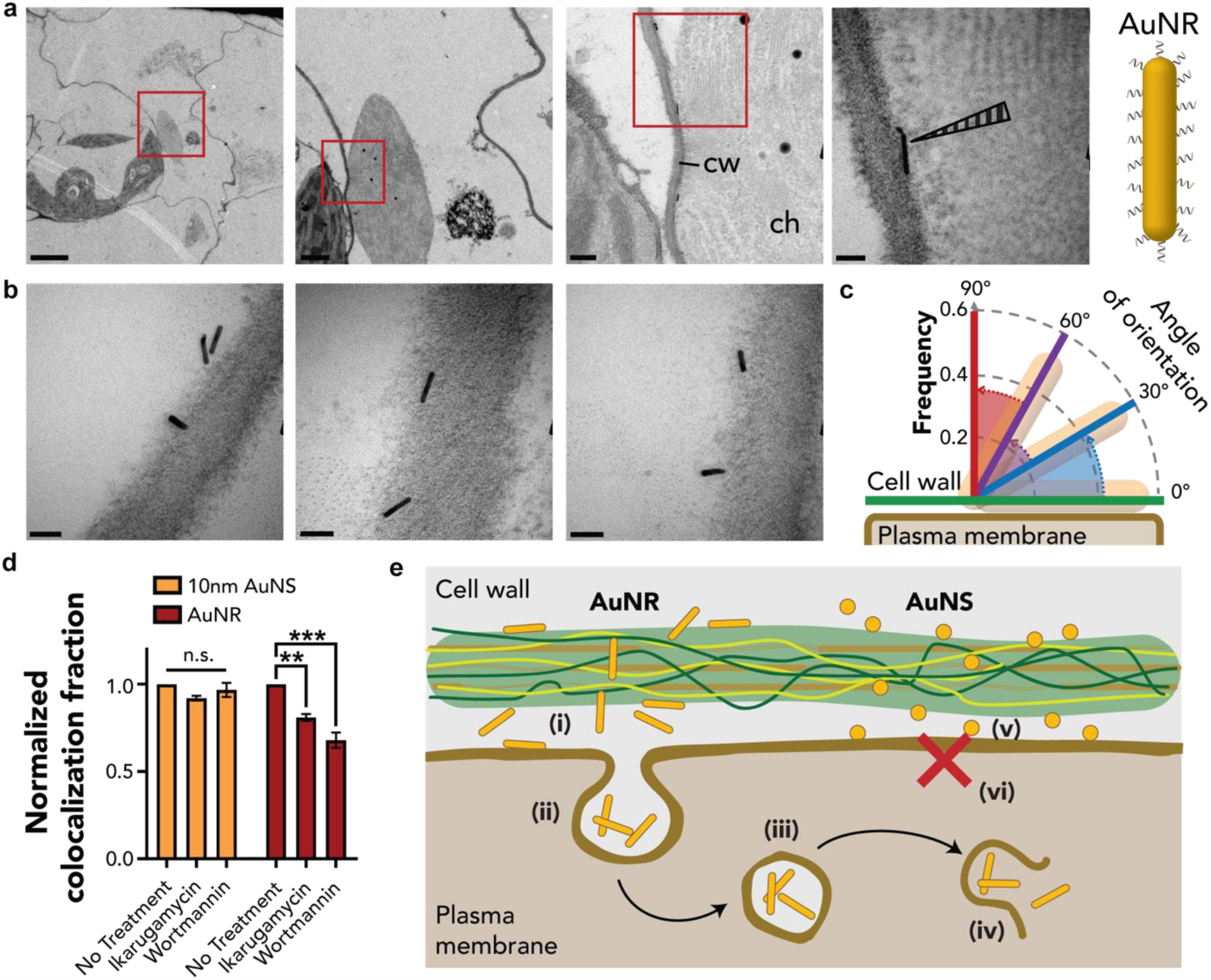
TEM of DNA-functionalized AuNR treated *Nicotiana benthamiana* leaves. **a** Representative TEM images for *Nb* plants 24h post-infiltration with DNA-functionalized AuNR. AuNR were identified within intact cells, suggesting AuNR can internalize into plant cells. Annotations represent cell wall (cw) and chloroplast (ch). Scale bars (from left to right) are 5 µm, 1 µm, 0.2 µm, and 50 nm. **b** AuNR exhibit a variety of orientations upon contact with the cell wall, with multiple observations of AuNR ‘piercing’ the cell wall, a potential mode of NP internalization. Scale bar: 50 nm. **c** Statistical analysis shows a dominant AuNR orientation that maximizes contact area with the cell wall (0° - 30°) and a low proportion of AuNR oriented perpendicular to the cell wall (60°-90°) for N = 324 AuNR. **d** Colocalization fractions between Cy3 and GFP channels for Cy3-tagged DNA-functionalized 10nm AuNS and AuNR infiltrated into *Nb* mGFP5 leaves treated with endocytosis inhibitors Ikarugamycin or Wortmannin. No change in colocalization fraction values are observed with AuNS, though AuNR samples in leaves treated with Ikarugamycin or Wortmannin demonstrate a significant decrease in colocalization fractions normalized to untreated samples, suggesting endocytosis is a mechanism for AuNR internalization. (AuNR, **p=0.007; ***p=0.0005 in two-way ANOVA, n.s.: not significant; error bars indicate s.e.m. n = 3). **e** Schematic of a potential mechanism for AuNS and AuNR association or internalization into plant cells. (i) AuNR exhibit various orientations relative to the cell wall, and may orient on their long ends to ‘pierce’ cell walls. (ii)-(iv) AuNR found in intact plant cells likely enter through endocytosis. (v) AuNS show high association with the cell wall, but (vi) no instances of any sized AuNS were identified inside plant cells within 24h.

Hypothesized mechanisms for NP internalization into plant cells include endocytosis, entry through the plasmodesmata, and NP acting as ‘nanospears’ to ‘pierce’ membranes^45^.To better understand how AuNR might internalize into plant cells, we assessed the impact of endocytosis inhibition on Cy3-DNA-AuNP association with plant cells using confocal microscopy. We infiltrated the leaves of GFP-expressing *Nb* with an endocytosis inhibitor (wortmannin or ikarugamycin), then infiltrated the same area with Cy3-DNA-AuNP (10 nm AuNS or AuNR) 30 minutes later. Infiltrated segments of the leaves were mounted and imaged with confocal microscopy 6 hours post-AuNP infiltration when colocalization levels above background signal could be observed, and colocalization analysis was performed by normalizing values for samples treated with endocytosis inhibitors to controls without inhibitors applied.

As shown in **Fig. 4d**, samples pre-treated with either endocytosis inhibitor induced a marked decrease in colocalization fraction for AuNR but no significant change for 10 nm AuNS (representative images in **Fig. S13**). The use of two types of endocytosis inhibitors, wortmannin, and ikarugamycin, yielded the same effects in both the 10 nm AuNS and AuNR. Representative images in **Fig. S13** showed that the colocalization of 10nm Cy3-DNA-AuNS was not affected by inhibitor treatment whereas the Cy3-DNA-AuNR showed a significant decrease in colocalization after treatment with either inhibitor. Wortmannin is an inhibitor of phosphoinositol-3 kinase activity and disrupts protein transport to vacuoles^46,47^. While the mechanism of action of ikarugamycin is undetermined, it is often used as an inhibitor of clathrin-mediated endocytosis^48,49^. Despite their different modes of action on plant cell trafficking, both wortmannin and ikarugamycin results suggest that a disruption of the plant cell’s native endocytic pathways result in different fates of sphere- and rod-shaped DNA-AuNPs upon association with plant cells. Specifically, we observe that endocytosis inhibition prevents AuNR entry into plant cells. These results are supported by TEM observations that AuNR internalize into plant cells, but not AuNS, thus we expect endocytosis inhibitors will not affect the association of 10 nm AuNS with the plant cell wall. Furthermore, our results suggest that endocytosis may be a mechanism for AuNR internalization into plant cells. Taken together, our TEM findings and endocytosis inhibitor assays suggest that AuNR internalize into plant cells through energy-dependent mechanisms, and AuNP do not internalize into plant cells. Based on these findings, we assembled a mechanism of AuNP transport within a plant leaf (**Fig. 4e**). Briefly, upon introduction to mature plant leaves, both DNA-AuNS and DNA-AuNR transport to the periphery of plant cells and experience a high degree of association with the cell wall. AuNR preferentially orient in parallel during initial contact with the cell wall and reorient perpendicularly when bypassing the cell wall, which supports proposed mechanisms of rotation-based cell wall entry of rod-shaped nanomaterials. After passage through the cell wall, endocytosis is a likely mechanism of AuNR cellular internalization.

### Sub-20 nm AuNP enable delivery of siRNA to silence mGFP5 in transgenic GFP *Nb* plant leaves

Targeted downregulation of certain plant proteins is known to confer disease resistance in crops. Therefore, an attractive alternative to the use of herbicides in agriculture is the use of a ‘green’ alternative known as RNA interference, which involves the delivery of small interfering RNA molecules (siRNA) that interfere with the production of a specific plant protein via sequence-specific destruction of the plant protein’s messenger RNA. However, siRNA is a molecule that is difficult to deliver across the lignin-rich and multi-layered cell wall as it is also highly susceptible to enzymatic degradation^50,51^. We therefore evaluated the use of AuNP as delivery vehicles to enable transient gene silencing through the delivery of siRNA into mature *Nb* plant leaves. Utilizing the low pH functionalization method, we synthesized all five AuNPs of varying morphologies and sizes with pre-hybridized siRNA targeting the GFP gene to obtain siRNA-functionalized AuNPs (siRNA-AuNPs, **Table S1**). In addition to DLS characterization, negative uranium staining of siRNA-AuNPs under TEM showed a visible halo surrounding AuNPs, suggesting successful siRNA-functionalization of the AuNPs (**Fig. 5a**, see **Fig. S1** for UV-Vis characterization and **Table S3** and **Fig. S14** for DLS characterization).

**Fig. 5.**
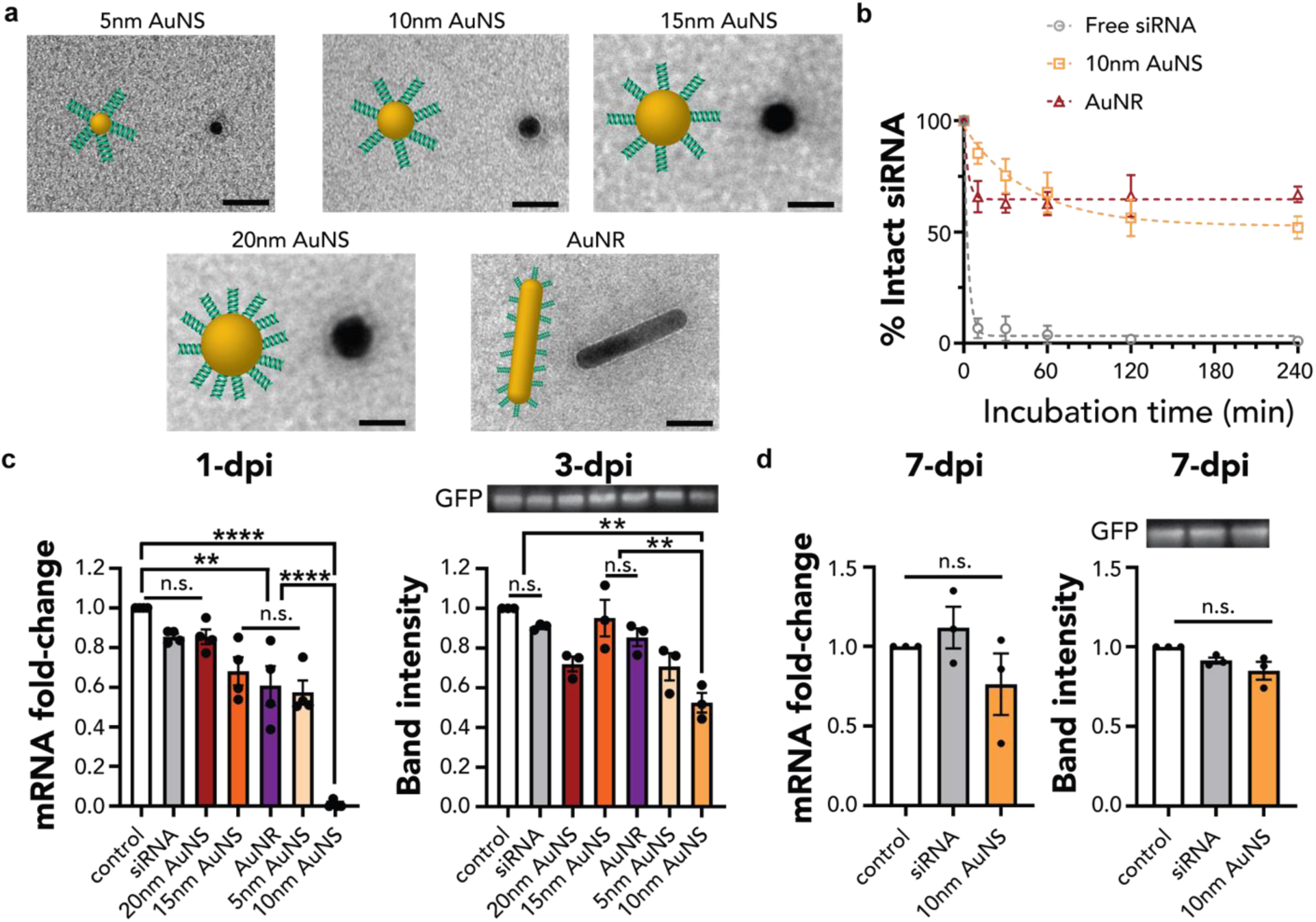
siRNA-functionalized AuNP exhibit size and morphology-dependent siRNA delivery efficiencies in mGFP5 *Nb* leaves. **a** TEM of stained siRNA-functionalized AuNP. The visible corona layer coating the AuNP confirms functionalization of AuNP with siRNA. Scale bar: 20nm. **b** AuNP protect siRNA from endoribonuclease RNase A degradation. siRNA-loaded 10nm AuNS and AuNR were incubated with RNase A for 10, 30, 60, 120, and 240 minutes prior to quantification of intact siRNA. Error bars indicate s.e.m. (n=3). **c** qPCR and Western blot analysis quantifying GFP mRNA fold-change and protein expression, 1 and 3 days respectively post-infiltration of mGFP5 *Nb* plant leaves with siRNA-AuNP. (qPCR: **p=0.0057, and ****p< 0.0001 in one-way ANOVA on day 1; n.s.: not significant; error bars indicate s.e.m. (n = 4).) (Western: **p=0.0020, 0.0074 in one-way ANOVA on day 3; error bars indicate s.e.m. (n=3).) **d** qPCR and Western blot analyses showing GFP mRNA and protein expression levels, respectively, return to baseline 7 days post-infiltration with siRNA-AuNP in *Nb* leaves. (qPCR: One-way ANOVA on day 7; n.s.: not significant; error bars indicate s.e.m. (n = 3).) (Western: One-way ANOVA on day 7; n.s.: not significant; error bars indicate s.e.m. (n=3).)

Prior to siRNA delivery in plants, we conducted an *in vitro* experiment to probe the ability of AuNP to protect siRNA from degradation. The density of siRNA per AuNP was determined using the aforementioned KCN-based decomposition method (**Table S5**). Next, siRNA-functionalized 10 nm AuNS and AuNR were incubated with a very high 1.2 μg/mL concentration of RNase A, to expedite the assay, for 0, 10, 30, 60, 120, and 240 minutes at room temperature. Following incubation, nuclease inactivation, and siRNA liberation from the AuNP surface, we quantified the concentration of remaining siRNA with a fluorescent plate reader. While free siRNA demonstrated extreme susceptibility to endonuclease degradation, a strong protective effect was afforded by the 10 nm AuNS and AuNR (**Fig. 5b**, see **Table S6** for statistical analysis). Free siRNA is largely degraded within 10 minutes of endonuclease exposure (7% ± 4 % remaining), while 10 nm AuNS and AuNR exhibit 54 ± 5 % and 64 ± 6 % intact siRNA by 240-minutes.

GFP-targeting siRNA-AuNPs were next infiltrated into GFP-expressing *Nb* leaves at a normalized 100 nM siRNA concentration, as previously used^11^. We next extracted mRNA and protein from treated leaf tissue and quantified the level of GFP silencing at the mRNA level 1-day post-infiltration *via* qPCR, and GFP-level silencing 3-days post-infiltration *via* a Western blot analysis (**Fig. 5c**). We initially hypothesized that only AuNR would enable efficient siRNA delivery and GFP silencing, since only AuNR were found to enter plant cells within the timescale of our experiments. Surprisingly, we did observe GFP silencing with both AuNS and AuNR, where smaller AuNS are generally more effective at GFP silencing than larger AuNS: 10 nm AuNS showed the greatest amount of silencing on both the mRNA (99%) and protein (48%) levels. Reduction in GFP mRNA transcripts in leaves 1-day post-infiltration were 15% (free siRNA), 15% (20 nm AuNS), 32% (15 nm AuNS), 39% (AuNR), 42% (5 nm AuNS), and 99% (10 nm AuNS) (**Fig. 5c**). Similarly, we measure a 9%, 28%, 5%, 15%, 29%, and 48% reduction in levels of GFP on Day 3 *via* Western Blot analysis of leaves treated with the aforementioned AuNPs, respectively (**Fig. 5c**). We also confirmed that gene silencing was transient *via* RT-qPCR and Western blot analyses of infiltrated leaves 7-days post-infiltration (**Fig. 5d**), in which mRNA and protein levels return to baseline several days after siRNA-AuNP treatment. We confirmed that GFP gene silencing was specific to the GFP-targeting siRNA sequence; 1-day post-infiltration qPCR analysis of nonsense siRNA-functionalized AuNS showed no silencing of the GFP gene (**Fig. S15**). Furthermore, we confirmed that functionalized siRNA can be detached from the surface of AuNP in apoplastic fluid (**Fig. S16**). These results stand in contrast with the assumption that NP cellular internalization is required for biomolecular delivery, whereby our results suggest that the internalization of NPs is not necessary for efficient siRNA delivery and gene silencing in plants.

Lastly, we confirmed the biocompatibility of nucleic acid functionalized-AuNP constructs by performing a quantitative PCR (qPCR) analysis of respiratory burst oxidase homolog B (*NbrbohB*) stress gene upregulation (**Fig. S17**). Results showed that neither 10 nm AuNS nor AuNR caused any significant change in *NbrbohB* gene expression, suggesting that AuNP can serve as a biocompatible siRNA-delivery vehicle to plants.

## Conclusion

Nanotechnology offers numerous advantages for the delivery of biological cargoes in plants^10–12^, yet an incomplete understanding of nanoparticle transport within plant tissues hinders rational design and development of abiotic plant delivery strategies. While studies investigate translocation and distribution of NPs within plants, often to understand their environmental effects^31,52,53^, the uncharacterized role of NP physicochemical characteristics on *in-planta* transport and the accessibility of their cargoes has made it difficult to develop delivery systems for fragile cargoes, such as siRNA, directly to plant cells.

Here, we systematically employed a series of DNA-AuNPs with various sizes (5-20 nm) and shapes (sphere and rod) to investigate how morphology impacts nanoparticle interaction with and uptake into plant cells following infiltration into mature *Nb* leaves. Confocal microscopy-based colocalization analysis demonstrated that smaller AuNS associate with plant cell walls faster than their larger counterparts. Our results confirm the plant cell wall size exclusion limit of ∼20 nm, whereby constructs of size exceeding the size exclusion limit cannot easily diffuse within plant tissues nor into plant cells. Prior literature in mammalian systems, such as simulations by Shi et al. and Jin et al., demonstrate that larger-sized AuNPs experience slower diffusion rates while increasing enthalpic contributions when interacting with surfaces that may encourage receptor-mediated endocytosis^54–56^. It is possible such size-dependent effects, albeit with a much more stringent size exclusion limit, are at play in plant cell NP transport.

NP-based bio-cargo delivery studies probing the uptake of NPs into plant cells rely on fluorescence microscopy^8–10^. Though optical imaging can visualize the presence and accumulation of fluorophore-tagged AuNP in the proximity of plant cells, resolution limitations prevent us from distinguishing more granular AuNP localization – AuNP could be associated with the outer cell wall, between the cell wall and cell membrane, or within the plant cell. Our findings from TEM microscopy, together with endocytosis inhibition assays, show that while sub-20 nm AuNS associate with plant cell walls, AuNS of all sizes are unlikely to enter plant cells, whereas AuNR do enter, likely through endocytosis. Furthermore, the 20 nm AuNS mainly found sandwiched between cell walls may be suggestive of their persistent fluorescent signal at long time scales but lower overall colocalization with the plant cell cytosol (**Fig. 2b**). Conversely, the 5, 10, and 15 nm AuNS experience greater freedom of movement and vulnerability to convective flows within leaf tissues, and are observed to accumulate on the extracellular side of individual cell walls. Analysis of hundreds of AuNRs suggest that rods may translocate across the plant cell wall through a rotation-mediated process that positions rods favorably at acute angles with respect to the plane of the cell wall; a mechanism that has been established for mammalian systems^21,23,57^. Theoretical and computational studies in mammalian systems have shown that the energetically favorable orientation of high-aspect ratio materials on lipid membranes is to increase the surface area of contact to the surface by lying flat. Although our surface of interest is the plant cell wall instead of cell membranes, our analysis shows similar trends as demonstrated by the higher than chance percentage of nanorods oriented between 0-10° to the cell wall. The internalization of AuNR and not AuNS is fascinating and stands in contrast to analogous size- and shape-based internalization studies performed in HeLa cells, where AuNS showed preferential internalization over AuNR, and in which 50 nm AuNS demonstrated highest levels of internalization^20^. We attribute these differences to the unique tissue morphologies of mature plant leaves, in particular the rigid cell wall. The theoretical underpinnings of how the plant cell wall influences plasma membrane behavior^58^ and subsequent internalization events remain under study, though it is known that the cell wall can regulate plasma membrane functions^59,60^. Our findings that internalization of AuNP into plant cells is shape-dependent motivates harnessing morphology in designing NP-based delivery tools for intracellular or extracellular delivery.

We further demonstrate that siRNA loaded AuNP below 20 nm in diameter, and AuNR, enable silencing of a GFP transgene in mature plants. Specifically, our study demonstrates that, within the timeframe of our experiments, AuNR can enter plant cells whereas AuNS likely do not, yet both enable siRNA delivery and gene silencing in mature plant leaves – with up to 99% gene silencing achieved by 10 nm siRNA-loaded AuNS. The greater efficiency of AuNP-based siRNA delivery relative to free siRNA delivery is likely due to AuNP-based protection of siRNA, whereby AuNP enable protection of siRNA against degradation for at least 1 day (**Fig. S18**). However, a more intriguing finding of our study is that AuNS enable siRNA delivery despite a lack of internalization into plant cells. We hypothesize that the interaction of the AuNP with individual plant cell walls increases their residence time proximal to cells, providing more opportunity for siRNA permeation and utilization into and by plant cells. Cell wall associated AuNP could act as a reservoir for siRNA, releasing siRNA from the gold surface upon association with biomolecules like thiol-containing compounds in the surrounding biofluids whilst simultaneously protecting the siRNA cargo from nuclease degradation. Furthermore, we found that 20 nm AuNP reside in between cells, but are not found imbedded inside cell walls – and that 20 nm AuNP are unable to deliver siRNA. These findings further suggest that while AuNP do not need to enter cells to deliver their cargoes, AuNP must at least reach the plant cell wall for their cargoes to be utilized: it is insufficient for AuNP to reside between cells for cargo delivery.

We therefore propose a mechanism of NP transport in plants in which NP morphology and size enable transport through plant leaf tissues, with various outcomes pertinent to cellular association and bio-cargo delivery. We highlight that smaller NP rather than larger NP translocate through plant tissues and localize in plant cell walls more efficiently. Interestingly, while rod-shaped NP enter plant cells, spherical NP do not, yet the ∼10 nm diameter of the latter is most efficient at accomplishing siRNA delivery through a proposed ‘reservoir’ mechanism in which NP imbed in the plant cell wall and protect siRNA from degradation while slowly releasing this cargo into plant cells for transient gene silencing. This study addresses a long-neglected gap between how NP transport in plant tissues is related to efficient delivery of surface-loaded biomolecular cargo and intracellular function. The unintuitive shape- and size-dependent transport, internalization, and silencing efficiencies of nucleic acid-functionalized AuNP motivate a non-canonical approach to development of future plant bionanotechnologies, including the rational design of AuNP for delivery of various payloads such as mRNA and protein to advance plant biotechnology and agriculture.

## Methods

### Chemicals and materials

Citrate-stabilized gold nanoparticles (AuNP), including gold nanospheres (5 nm, 10 nm, 15 nm and 20 nm) and gold nanorods (13 nm x 68 nm) were purchased from NanoComposix, and used for all AuNP-based experiments. Wortmannin (19545-26-7, 1mg) was purchased from FisherScientific, and ikarugamycin (36531-78-9, 500 μg) was purchased from Cayman Chemicals. SYBR Gold was purchased from Thermo Fisher. RNase A was purchased from Takara Bio. The following chemicals were purchased from Sigma-Aldrich: citric acid, sodium chloride, potassium chloride, sodium phosphate dibasic, sodium phosphate monobasic (anhydrous), Tris hydrochloride, glycerol, diethyl pyrocarbonate (DEPC), NP-40, and Protease Inhibitor Cocktail (for plant cell extracts). Ethylenediaminetetraacetic acid (EDTA) and Bovine Serum Albumin (BSA) were purchased from FisherScientific. The RNeasy plant mini kit was purchased from QIAGEN, iScript cDNA synthesis kit from Bio-Rad, PowerUp SYBR green master mix from Applied Biosystems, Pierce 660 nm assay and Pierce ECL Western Blotting Substrate from Thermo Fisher, pre-cast Tris/Glycine gels 4-20% gradient, 10x TBS buffer and 10x Tris/Glycine buffer from BIORAD, and antibodies from Abcam. Single stranded RNA and DNA oligonucleotides were purchased from and purified by Integrated DNA Technologies, Inc. (IDT); DNA oligonucleotides labelled with Cy3 were purified by HPLC, and all the oligonucleotides were dissolved in DNase/RNase-free water before use. The concentration of each strand was estimated using absorbance measurements at 260 nm and the corresponding extinction coefficients provided by IDT.

### Plant growth, Maintenance, and Leaf Infiltration

Transgenic mGFP5 *Nicotiana benthamiana* seeds (obtained from the Staskawicz Lab, UC Berkeley) were germinated and grown in SunGro Sunshine LC1 Grower soil mixture within the growth chamber (740 FHLED, HiPoint). These mGFP5 Nb plants encode for GFP with a ER-localization signal, yet confocal imaging (unpublished) demonstrates a qualitatively distinct profile from the commonly-used 16C Nb variant which is also GFP-expressing. Nevertheless, GFP signal from mGFP5 Nb originates from the cytosol, and this phenomenon is used for downstream analyses. The plants were grown in 4-inch pots under LED light (100 µmol/m^2^-s) and a 14/10 light/dark photoperiod at 23 °C and 60% humidity. All experiments were done with intact leaves in plants 4 weeks of age, where plants were incubated in the growth chamber until the time of data collection. Young leaves (approximately 2 x 3 cm in size) were infiltrated with a 1 ml needleless syringe on the abaxial side of the leaves after a tiny puncture had been made with a 10µL pipette tip.

### Nucleic acid (NA)-AuNP Functionalization

#### NA-functionalization of citrate-stabilized AuNP

AuNP were functionalized with thiol-functionalized DNA or RNA (SH-DNA/SH-RNA, see detailed sequences in table S1) by using a facile pH-assisted method reported previously by Zhang et al^40^. The pH-assisted method generates functional DNA-functionalized AuNP (DNA-AuNP) constructs in a matter of hours, as opposed to the salt-aging method whereby the AuNP salt concentration is progressively increased by the stepwise addition of salt over several hours and often requires an overnight incubation. AuNP were concentrated in a refrigerated centrifuge (5424R, Eppendorf) to their desired concentration, resuspended, then incubated with SH-DNA or SH-RNA (SH-NA) at specific ratios (see details in table S2) for 5 minutes. The pH of the solution was lowered by adding 500 mM citrate buffer (pH 3) for a final concentration of 10 mM, followed by pipette mixing. After 15 minutes, the pH of the solution was adjusted to neutral with 200 mM PB buffer (pH 7.6) for a final concentration of 10 mM and the solution left to sit for 15 minutes. To remove free SH-NA, samples were centrifuged and washed three times with 10 mM PB buffer (pH 7.6) whereby free SH-NA could no longer be detected in the supernatant. The nanoparticles were resuspended in 0.3x PBS to obtain DNA- or RNA-functionalized AuNP at desired concentrations.

#### Fluorophore-tagged NA-AuNP

Fluorophore-tagged AuNP (Cy3-DNA-AuNP) were obtained whereby Cy3-DNA with a 15-nt sequence complementary to DNA (Table S1) on the AuNP was added to DNA-AuNP in 0.3x PBS (41 mM NaCl, 0.8 mM KCl, 2.4 mM Na2HPO4, and 0.6 mM KH2PO4). The Cy3-DNA-AuNP solution was then incubated at 40°C for 30 min, then cooled to room temperature for at least an hour prior to use.

### Nanoparticle Characterization

#### UV-Vis-IR spectroscopy measurements

100 µL of AuNP and DNA or RNA modified AuNP in 0.3x PBS were added in Sub-Micro Quartz Cuvettes (Cole-Parmer), and the spectrophotometric measurements were carried out with a UV-3600 Plus UV-Vis-NIR Spectrophotometer (Shimadzu North America).

#### Dynamic light scattering (DLS) measurements

DLS measurements were taken with the Zetasizer Nano ZS (Malvern Analytical). Citrate-stabilized AuNP and NA-AuNP in 2mM citrate and 0.3x PBS buffer respectively were diluted to OD1, placed in disposable cuvettes (Malvern), and set up for DLS measurement.

#### Transmission electron microscopy (TEM)

The structure of the AuNP was examined using a JEOL 1200EX instrument, and the interactions of AuNP with plant cells was studied with a Tecnai 12 TEM (Berkeley Electron Microscope Lab). AuNP were drop casted on plasma-treated Formvar carbon-coated TEM grids (Ted-Pella), allowed to sit for 10 minutes, then the remaining solution wicked off before imaging at an acceleration voltage of 120 kV. To visualize siRNA on NA-AuNP, TEM grids with drop casted NA-AuNP were placed face-down on a 2% methanolic uranyl acetate droplet for 30 seconds, removed, and blotted with filter paper and air-dried prior to imaging.

#### Quantification of DNA on DNA-AuNP

The number of DNA strands on each functionalized gold nanoparticle was quantified via a KCN desorption assay previously reported by Baldock et al^42^. 100mM KCN solution (pH = 12) was added to a known concentration (calculated from the UV-Vis-IR spectra) of DNA-AuNP. After being left overnight, the absorbance spectrum was collected, and Beer’s Law used to calculate DNA concentration at 260nm. The number of DNA strands per AuNP was calculated by dividing the DNA concentration by the AuNP concentration.

### Tracking of AuNP in *Nb* leaves

#### Co-localization analysis of Cy3 labelled AuNP with GFP

100uL Cy3-DNA-AuNP at a Cy3 concentration of 400 nM were infiltrated into plant leaves and left on the benchtop at 20°C for the desired incubation time. For samples used to investigate nanoparticle internalization pathway, 40 μM wortmannin or 10 μM ikarugamycin solutions were infiltrated into target leaves 30 minutes prior to Cy3-DNA-AuNP introduction. To prepare infiltrated leaves for confocal imaging, a small leaf section was cut and mounted between a glass slide and #1 thickness cover slip. Water was added to keep the leaf sections hydrated during imaging. A Zeiss LSM 710 confocal microscope was used to image the plant tissue with 488 nm and 543 nm laser excitation for GFP and Cy3 signal collection respectively. The collection window for Cy3 was adjusted to 550 – 604 nm to avoid crosstalk between Cy3 and leaf chlorophyll autofluorescence. Images were obtained at 20x magnification. The same imaging parameters and quantification analyses were applied to samples imaged on different days.

#### Transmission Electron Microscopy

For internalization study of AuNP, leaves infiltrated with AuNP were cut into small pieces approximately 1mm by 3mm in size. Leaf samples were fixed using 2% glutaraldehyde in 0.1 M sodium cacodylate buffer (pH 7.2), then subject to microwave-assisted vacuum to remove air in the vacuoles. Samples were post-fixed with 1% osmium tetroxide in 0.1 M sodium cacodylate buffer (pH 7.2), dehydrated with acetone, and transferred into epoxy resin for embedding. Finally, epoxy resin-embedded samples were cut into 100-nm-thin cross-sectioned films using a Reichert-Jung Ultracut E microtome, then transferred onto bare Cu TEM grids for imaging at an acceleration voltage of 120 kV.

### Nuclease Protection

To demonstrate the protective effect of AuNP on cargo, a nuclease protection assay was carried out. siRNA functionalized AuNP (concentrations corresponding to approximately 0.2µM siRNA) were incubated with RNase A (final concentration 1.2 µg/mL) at room temperature for 0, 10, 30, 60, 120, and 240 minutes. Post-incubation, the endonuclease was inactivated with DEPC addition, and KCN added to a final concentration of 18mM to facilitate NP decomposition. Post-overnight incubation at 4°C, 20µL of the samples were added to 180uL of Quant-iT microRNA Assay Kit (Life Technologies, Thermo Fisher Scientific) reagents and fluorescence was read with an Infinite M1000 PRO microplate reader (Tecan). Free siRNA controls were included for each timepoint. Duplicate measurements were made for each of the three experimental replicates. In a separate experiment, DNA-AuNP loaded with siRNA containing at 15-nt complementary overhang were incubated with RNase A at room temperature for 0, 1, 6, 12, 18, and 24h. Post-incubation, the nuclease was inactivated with DEPC addition, then the AuNP solution treated with 8M urea at 37°C for 30min to disrupt interactions between DNA AuNP and siRNA. Free siRNA controls were included for each timepoint. Samples were loaded and run on a 3% agarose gel pre-stained with SYBR Gold at 70V for 25min. The resulting gels were imaged using a Typhoon FLA 9500 (General Electric), and gel band intensities analyzed using FIJI and GelBandFitter^61^.

### Analyzing GFP Silencing in Nb leaves using siRNA-AuNP

#### Reverse Transcriptase-Quantitative PCR

siRNA-AuNP loaded with 100 nM siRNA were infiltrated into plant leaves and left on the benchtop at 20°C for 24 hours or 7 days upon which total RNA was extracted. Two-step qPCR was performed to quantify GFP gene silencing with the following commercially available kits: RNeasy plant mini kit (QIAGEN) for total RNA extraction from leaves, iScript cDNA synthesis kit (Bio-Rad) to reverse transcribe total RNA into cDNA, and PowerUp SYBR green master mix (Applied Biosystems) for qPCR. The target gene in our qPCR was mGFP5 (GFP transgene inserted into *Nicotiana benthamiana*), and EF-1 (elongation factor 1) was chosen as the housekeeping (reference) gene^62,63^. Primers (see detailed sequences in Table S1) for these genes (fGFP, rGFP, fEF1 and rEF1) were ordered from IDT and used without further purification. An annealing temperature of 60°C was used for qPCR, which was run for 40 cycles. qPCR data was analyzed by the ddCt method^64^ to obtain the normalized GFP gene expression-fold change with respect to the EF-1 housekeeping gene and control sample. For each sample, qPCR was performed as 3 technical replicates (3 reactions from the same isolated RNA batch), and the entire experiment consisting of independent infiltrations and RNA extractions from different plants was repeated 3 times (3 biological replicates).

#### Western Blot

siRNA-AuNP loaded with 100 nM siRNA were infiltrated into plant leaves and left on the benchtop at 20°C for 2 or 7 days upon which proteins were extracted. Briefly, infiltrated leaves were cut, frozen in liquid nitrogen, and ground with a mortar and pestle. 350uL lysis buffer (10 mM Tris/HCl (pH 7.5), 150 mM NaCl, 1 mM EDTA, 0.1% NP-40, 5% glycerol, and 1% protease inhibitor cocktail) was added to the resulting powder, vortexed, spun down, and kept on ice. All samples were incubated at 50°C for 3 minutes, centrifuged at 16,000g for 30 minutes, and the supernatant transferred to a new tube to obtain extracted proteins. Protein concentration of samples were quantified using a Pierce 660 nm Protein Assay (Thermo Fisher), and concentrations of proteins standardized with the addition of lysis buffer. Loading dye was added to protein solutions, incubated at 95°C for 10 minutes, and centrifuged at 16,000g for 15 minutes. A 4-20% Mini-PROTEAN Precast Protein Gel (BIORAD) was loaded with samples and run at 120V for 60 minutes with a Mini-PROTEAN Tetra Cell (BIORAD). The gel was included in a sandwich with a methanol activated PVDF membrane, placed in cold 1x Tris/Glycine buffer, and run at 400 mA for 60 minutes. Post-transfer, the membrane was rinsed with 1x TBST buffer three times, with 5 minutes in between each rinse, followed by a 60-minute incubation with 5% BSA in TBST, and three rinses with 1x TBST including incubations. The membrane was incubated at 4°C on a shaker with rabbit antibody anti-GFP (1:3000 dilution) overnight, rinsed with TBST, incubated for 60 minutes with goat anti-rabbit antibody (1:5000 dilution), rinsed with TBST. The membrane was washed briefly in MilliQ water, exposed to Pierce ECL Western Blotting Substrate (Thermo Scientific), and immediately imaged. 3 biological replicates were collected, and band intensity analyses done on each.

### siRNA Desorption in Plant Biofluid

Apoplastic fluid from mature leaves was extracted from month-old Nb plants following the protocol outlined by O’Leary et al^65^. Briefly, 4-5 Nb leaves were detached at the petiole, washed in MilliQ water, rolled up and placed into a 60 mL syringe with a syringe cap. The syringe was filled with MilliQ water to the 40 mL mark, and air expelled from the syringe. The plunger was pulled slowly to the 60 mL mark to induce negative pressure, slowly released back to 40 mL, and excess air ejected. This vacuum infiltration was performed three times, upon which the leaves were removed from the syringe and gently patted dry. Next, a 1 mL pipette tip was placed at the end of a strip of Parafilm (∼5 in) and leaves oriented with petioles on one side. The Parafilm was rolled up into a 20 mL syringe (no plunger) with petioles facing up, and the 20 mL syringe put into a 50 mL Falcon tube. The tube was centrifuged for 10 min at 1,000x rpm at 4°C with minimum acceleration and deceleration (5810R Centrifuge, Eppendorf). Any fluid expelled was collected into 1.5 mL tubes, centrifuged at 15,000x rpm for 5 minutes, and the supernatant collected to obtain apoplastic fluid. Plant lysate was obtained in an identical fashion to protein extraction for Western Blot analysis as described previously. The protein content of the apoplastic fluid and lysate was quantified using the Pierce 660 nm Protein Assay.

2 ug of protein was incubated with 10 nm siRNA-AuNP (final concentration of 300 nM siRNA) on the benchtop for 24 hours, upon which the AuNP were centrifuged and the supernatant collected. The supernatant was run on a 3% agarose gel to visualize any intact siRNA that had desorbed from the siRNA-AuNP.

### Statistics and Data Analysis

#### Confocal Cy3-GFP co-localization data

Data collection for each sample was done in triplicate, with 10-15 technical replicates (non-overlapping confocal field of views from each leaf) collected for each biological replicate (an infiltration into a unique plant). Each field of view was analyzed with ImageJ analysis software to obtain a co-localization proportion as given by the Mander’s Overlap Coefficient, and all fields of view were averaged to obtain a single value for that biological replicate. Data are expressed as each mean from the 3 biological replicates, with error bars denoting standard error of the mean. Significance is measured with one-way ANOVA with Tukey’s multiple comparisons test. For the incubation time-varied experiments, in order of the 5 nm AuNS, 10 nm AuNS, 15 nm AuNS, 20 nm AuNS, and AuNR confocal experiments, F = 36.07, 21.95, 18.65, 35.29, and 75.64, where the corresponding P-value for all the aforementioned experiments was **** P<0.0001. For the endocytosis inhibitor assay, the significance of the 10 nm AuNS and AuNR results was similarly measured with one-way ANOVA with Tukey’s multiple comparisons test; F = 2.621 and P = 0.1520, F = 33.46 and P = 0.0006 (***) respectively.

#### TEM data

To quantify angle of NR orientation with respect to the cell wall, ImageJ’s Measure function was used to draw tangents to local cell wall contacting NR and direction of NR long axis. The resulting difference in angle (acute) was used.

#### Nuclease protection data

For Quant-iT based quantification, data collection for each sample was done in triplicate, with fluorescence measurements performed in duplicate and the numerical average used for further analysis. Background fluorescence from remnant DNA and background noise was subtracted from all samples. The proportion of intact siRNA at time T was obtained using the following equation,

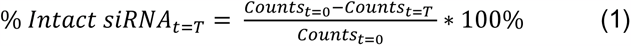

Data are expressed as each mean from the 3 replicates, with error bars denoting standard error of the mean. Significance is measured with two-way ANOVA with Tukey’s multiple comparisons test; for time (n = 6), column factor (n = 3), and subject (n = 9), F = 267.6 and P<0.0001, F = 186.7 and P<0.0001, and F = 4.953 and P = 0.0013 respectively.

For the gel-based quantification, data collection for each sample was done in triplicate. Following band intensity quantification and deconvolution, the proportion of intact siRNA at time T was obtained using an analogous equation to Equation (1). Data are expressed as each mean from the 3 replicates, with error bars denoting standard deviation. Significance is measured with two-way ANOVA with Tukey’s multiple comparisons test; for time (n = 5), column factor (n = 3), and subject (n = 9), F = 21.07 and P = 0.0010, F = 45.85 and P = 0.0002, and F = 26.11 and P<0.0001 respectively.

#### qPCR data

GFP silencing experiments comprised of 3 biological replicates, whereby 3 separate leaves are infiltrated per sample and analyzed with qPCR. Each biological replicate consisted of 3 technical replicates for the qPCR reaction. No template controls and no reverse transcriptase controls were also performed. Data are expressed as each mean from the 3 biological replicates, with error bars denoting standard error of the mean. Significance is measured with one-way ANOVA with Tukey’s multiple comparisons test. For 1-day and 7-day experiments, F = 36.76 and P<0.0001, F = 1.803 and P = 0.2436 respectively.

#### Western blot data

Western blot experiments to quantify GFP comprised of 3 biological replicates, whereby 3 separate leaves are infiltrated per sample and proteins extracted for further analysis. Data are expressed as the mean from the 3 biological replicates, with error bars denoting standard error of the mean. Significance is measured with one-way ANOVA with Tukey’s multiple comparisons test. For 3-day and 7-day experiments, F = 10.26 and P = 0.0002, F = 2.122 and P = 0.2010 respectively.

## Acknowledgements

We acknowledge support of a Burroughs Wellcome Fund Career Award at the Scientific Interface (CASI), a Stanley Fahn PDF Junior Faculty Grant with Award # PF-JFA-1760, a Beckman Foundation Young Investigator Award, a USDA AFRI award, a USDA NIFA award, and a Foundation for Food and Agriculture Research (FFAR) New Innovator Award (M.P.L). M.P.L. is a Chan Zuckerberg Biohub investigator. N.S.G. is supported by a FFAR Fellowship. J.W.W. is a receipient of the National Science Foundation Graduate Research Fellowship. S-J.P. acknowledges support from LG Yonam Foundation. We acknowledge the support of UC Berkeley CRL Molecular Imaging Center, the UC Berkeley Electron Microscopy Lab, and the Innovative Genomics Institute (IGI). Work at the Molecular Foundry was supported by the Office of Science, Office of Basic Energy Sciences, of the U.S. Department of Energy under Contract No. DE-AC02-05CH11231.

## Author Contributions

H.Z. and N.S.G. conceived of the project, designed the study, and wrote the manuscript. H.Z. and N.S.G. performed the majority of experiments and data analysis. J.W.W. and G.S.D. contributed key input and advanced project direction. S.B. performed initial experiments verifying project feasibility. All authors have edited and commented on the manuscript and have given their approval of the final version.

**Correspondence and requests for materials** should be addressed to M.P.L.

## References

1. Farokhzad, O. C. & Langer, R. Nanomedicine: developing smarter therapeutic and diagnostic modalities. Adv. Drug Deliv. Rev. 58, 1456–1459 (2006).

2. Parveen, S., Misra, R. & Sahoo, S. K. Nanoparticles: a boon to drug delivery, therapeutics, diagnostics and imaging. Nanomedicine Nanotechnology, Biol. Med. 8, 147–166 (2012).

3. Chen, G., Roy, I., Yang, C. & Prasad, P. N. Nanochemistry and Nanomedicine for Nanoparticle-based Diagnostics and Therapy. Chem. Rev. 116, 2826–2885 (2016).

4. Liu, Y. et al. A gene cluster encoding lectin receptor kinases confers broad-spectrum and durable insect resistance in rice. Nat. Biotechnol. 33, 301–305 (2015).

5. Li, T., Liu, B., Spalding, M. H., Weeks, D. P. & Yang, B. High-efficiency TALEN-based gene editing produces disease-resistant rice. Nature biotechnology 30, 390–392 (2012).

6. Karny, A., Zinger, A., Kajal, A., Shainsky-Roitman, J. & Schroeder, A. Therapeutic nanoparticles penetrate leaves and deliver nutrients to agricultural crops. Sci. Rep. 8, 7589 (2018).

7. Torney, F., Trewyn, B. G., Lin, V. S. Y. & Wang, K. Mesoporous silica nanoparticles deliver DNA and chemicals into plants. Nat. Nanotechnol. 2, 295– 300 (2007).

8. Demirer, G. S. et al. High aspect ratio nanomaterials enable delivery of functional genetic material without DNA integration in mature plants. Nat. Nanotechnol. 14, 456–464 (2019).

9. Kwak, S.-Y. et al. Chloroplast-selective gene delivery and expression in planta using chitosan-complexed single-walled carbon nanotube carriers. Nat. Nanotechnol. 14, 447–455 (2019).

10. Mitter, N. et al. Clay nanosheets for topical delivery of RNAi for sustained protection against plant viruses. Nat. Plants 3, 16207 (2017).

11. Demirer, G. S. et al. Carbon nanocarriers deliver siRNA to intact plant cells for efficient gene knockdown. Sci. Adv. 6, eaaz0495 (2020).

12. Zhang, H. et al. DNA nanostructures coordinate gene silencing in mature plants. Proc. Natl. Acad. Sci. 116, 7543 LP – 7548 (2019).

13. Martin-Ortigosa, S. et al. Mesoporous silica nanoparticle-mediated intracellular cre protein delivery for maize genome editing via loxP site excision. Plant Physiol. 164, 537–47 (2014).

14. Liu, Q. et al. Carbon Nanotubes as Molecular Transporters for Walled Plant Cells. Nano Lett. 9, 1007–1010 (2009).

15. Bao, W., Wang, J., Wang, Q., O’Hare, D. & Wan, Y. Layered Double Hydroxide Nanotransporter for Molecule Delivery to Intact Plant Cells. Sci. Rep. 6, 26738 (2016).

16. Wong, M. H. et al. Lipid Exchange Envelope Penetration (LEEP) of Nanoparticles for Plant Engineering: A Universal Localization Mechanism. Nano Lett. 16, 1161– 1172 (2016).

17. Zhang, S., Gao, H. & Bao, G. Physical Principles of Nanoparticle Cellular Endocytosis. ACS Nano 9, 8655–8671 (2015).

18. Herd, H. et al. Nanoparticle Geometry and Surface Orientation Influence Mode of Cellular Uptake. ACS Nano 7, 1961–1973 (2013).

19. Xie, X., Liao, J., Shao, X., Li, Q. & Lin, Y. The Effect of shape on Cellular Uptake of Gold Nanoparticles in the forms of Stars, Rods, and Triangles. Sci. Rep. 7, 3827 (2017).

20. Chithrani, B. D., Ghazani, A. A. & Chan, W. C. W. Determining the Size and Shape Dependence of Gold Nanoparticle Uptake into Mammalian Cells. Nano Lett. 6, 662–668 (2006).

21. Yi, X., Shi, X. & Gao, H. A Universal Law for Cell Uptake of One-Dimensional Nanomaterials. Nano Lett. 14, 1049–1055 (2014).

22. Shi, X., von dem Bussche, A., Hurt, R. H., Kane, A. B. & Gao, H. Cell entry of one-dimensional nanomaterials occurs by tip recognition and rotation. Nat. Nanotechnol. 6, 714–719 (2011).

23. Huang, C., Zhang, Y., Yuan, H., Gao, H. & Zhang, S. Role of Nanoparticle Geometry in Endocytosis: Laying Down to Stand Up. Nano Lett. 13, 4546–4550 (2013).

24. Vácha, R., Martinez-Veracoechea, F. J. & Frenkel, D. Receptor-Mediated Endocytosis of Nanoparticles of Various Shapes. Nano Lett. 11, 5391–5395 (2011).

25. Hui, Y. et al. Role of Nanoparticle Mechanical Properties in Cancer Drug Delivery. ACS Nano 13, 7410–7424 (2019).

26. Houston, K., Tucker, M. R., Chowdhury, J., Shirley, N. & Little, A. The Plant Cell Wall: A Complex and Dynamic Structure As Revealed by the Responses of Genes under Stress Conditions. Frontiers in Plant Science 7, 984 (2016).

27. Cunningham, F. J., Goh, N. S., Demirer, G. S., Matos, J. L. & Landry, M. P. Nanoparticle-Mediated Delivery towards Advancing Plant Genetic Engineering. Trends Biotechnol. 36, 882–897 (2018).

28. Schwab, F. et al. Barriers, pathways and processes for uptake, translocation and accumulation of nanomaterials in plants – Critical review. Nanotoxicology 10, 257–278 (2016).

29. Wang, P., Lombi, E., Zhao, F. J. & Kopittke, P. M. Nanotechnology: A New Opportunity in Plant Sciences. Trends in Plant Science 21, 699–712 (Elsevier Current Trends, 2016).

30. Hubbard, J. D., Lui, A. & Landry, M. P. Multiscale and multidisciplinary approach to understanding nanoparticle transport in plants. Curr. Opin. Chem. Eng. 30, 135–143 (2020).

31. Corredor, E. et al. Nanoparticle penetration and transport in living pumpkin plants: in situsubcellular identification. BMC Plant Biol. 9, 45 (2009).

32. Bao, D., Oh, Z. G. & Chen, Z. Characterization of Silver Nanoparticles Internalized by Arabidopsis Plants Using Single Particle ICP-MS Analysis. Frontiers in Plant Science 7, 32 (2016).

33. Avellan, A. et al. Nanoparticle Size and Coating Chemistry Control Foliar Uptake Pathways, Translocation, and Leaf-to-Rhizosphere Transport in Wheat. ACS Nano 13, 5291–5305 (2019).

34. Zhang, P. et al. Shape-Dependent Transformation and Translocation of Ceria Nanoparticles in Cucumber Plants. Environ. Sci. Technol. Lett. 4, 380–385 (2017).

35. Naha, P. C., Chhour, P. & Cormode, D. P. Systematic in vitro toxicological screening of gold nanoparticles designed for nanomedicine applications. Toxicol. Vitr. 29, 1445–1453 (2015).

36. Rana, S., Bajaj, A., Mout, R. & Rotello, V. M. Monolayer coated gold nanoparticles for delivery applications. Adv. Drug Deliv. Rev. 64, 200–216 (2012).

37. Daniel, M. C. & Astruc, D. Gold Nanoparticles: Assembly, Supramolecular Chemistry, Quantum-Size-Related Properties, and Applications Toward Biology, Catalysis, and Nanotechnology. Chem. Rev. 104, 293–346 (2004).

38. Pei, H. et al. Designed Diblock Oligonucleotide for the Synthesis of Spatially Isolated and Highly Hybridizable Functionalization of DNA–Gold Nanoparticle Nanoconjugates. J. Am. Chem. Soc. 134, 11876–11879 (2012).

39. Yao, G. et al. Programming nanoparticle valence bonds with single-stranded DNA encoders. Nat. Mater. 19, 781–788 (2020).

40. Zhang, X., Servos, M. R. & Liu, J. Instantaneous and Quantitative Functionalization of Gold Nanoparticles with Thiolated DNA Using a pH-Assisted and Surfactant-Free Route. J. Am. Chem. Soc. 134, 7266–7269 (2012).

41. Wei, G. et al. Hairpin DNA-functionalized gold nanorods for mRNA detection in homogenous solution. J. Biomed. Opt. 21, 1–9 (2016).

42. Baldock, B. L. & Hutchison, J. E. UV–Visible Spectroscopy-Based Quantification of Unlabeled DNA Bound to Gold Nanoparticles. Anal. Chem. 88, 12072–12080 (2016).

43. Geilfus, C.-M. & Muehling, K. Real-Time Imaging of Leaf Apoplastic pH Dynamics in Response to NaCl Stress. Frontiers in Plant Science 2, 13 (2011).

44. Yu, M. et al. Rotation-Facilitated Rapid Transport of Nanorods in Mucosal Tissues. Nano Lett. 16, 7176–7182 (2016).

45. Foroozandeh, P. & Aziz, A. A. Insight into Cellular Uptake and Intracellular Trafficking of Nanoparticles. Nanoscale Res. Lett. 13, 339 (2018).

46. Matsuoka, K., Bassham, D. C., Raikhel, N. V & Nakamura, K. Different sensitivity to wortmannin of two vacuolar sorting signals indicates the presence of distinct sorting machineries in tobacco cells. J. Cell Biol. 130, 1307–1318 (1995).

47. Aniento, F. & Robinson, D. G. Testing for endocytosis in plants. Protoplasma 226, 3–11 (2005).

48. Reynolds, G. D., Wang, C., Pan, J. & Bednarek, S. Y. Inroads into Internalization: Five Years of Endocytic Exploration. Plant Physiol. 176, 208–218 (2018).

49. Elkin, S. R. et al. Ikarugamycin: A Natural Product Inhibitor of Clathrin-Mediated Endocytosis. Traffic 17, 1139–1149 (2016).

50. Meister, G. & Tuschl, T. Mechanisms of gene silencing by double-stranded RNA. Nature 431, 343–349 (2004).

51. Tiwari, M., Sharma, D. & Trivedi, P. K. Artificial microRNA mediated gene silencing in plants: progress and perspectives. Plant Mol. Biol. 86, 1–18 (2014).

52. Zhu, Z.-J. et al. Effect of Surface Charge on the Uptake and Distribution of Gold Nanoparticles in Four Plant Species. Environ. Sci. Technol. 46, 12391–12398 (2012).

53. Zhai, G., Walters, K. S., Peate, D. W., Alvarez, P. J. J. & Schnoor, J. L. Transport of Gold Nanoparticles through Plasmodesmata and Precipitation of Gold Ions in Woody Poplar. Environ. Sci. Technol. Lett. 1, (2014).

54. Chithrani, D. B. Intracellular uptake, transport, and processing of gold nanostructures. Mol. Membr. Biol. 27, 299–311 (2010).

55. Shi, W., Wang, J., Fan, X. & Gao, H. Size and shape effects on diffusion and absorption of colloidal particles near a partially absorbing sphere: Implications for uptake of nanoparticles in animal cells. Phys. Rev. E 78, 61914 (2008).

56. Jin, H., Heller, D. A., Sharma, R. & Strano, M. S. Size-Dependent Cellular Uptake and Expulsion of Single-Walled Carbon Nanotubes: Single Particle Tracking and a Generic Uptake Model for Nanoparticles. ACS Nano 3, 149–158 (2009).

57. Yang, H. et al. Mechanism for the Cellular Uptake of Targeted Gold Nanorods of Defined Aspect Ratios. Small 12, 5178–5189 (2016).

58. Cheddadi, I., Génard, M., Bertin, N. & Godin, C. Coupling water fluxes with cell wall mechanics in a multicellular model of plant development. PLOS Comput. Biol. 15, e1007121 (2019).

59. McKenna, J. F. et al. The cell wall regulates dynamics and size of plasma-membrane nanodomains in Arabidopsis Proc. Natl. Acad. Sci. 116, 12857 LP – 12862 (2019).

60. Liu, Z., Persson, S. & Sánchez-Rodríguez, C. At the border: the plasma membrane–cell wall continuum. J. Exp. Bot. 66, 1553–1563 (2015).

61. Mitov, M. I., Greaser, M. L. & Campbell, K. S. GelBandFitter – A computer program for analysis of closely spaced electrophoretic and immunoblotted bands. Electrophoresis 30, 848–851 (2009).

62. Nicot, N., Hausman, J.-F., Hoffmann, L. & Evers, D. Housekeeping gene selection for real-time RT-PCR normalization in potato during biotic and abiotic stress. J. Exp. Bot. 56, 2907–2914 (2005).

63. Selvakesavan, R. K. & Franklin, G. Nanoparticles Affect the Expression Stability of Housekeeping Genes in Plant Cells. Nanotechnol. Sci. Appl. Volume 13, 77–88 (2020).

64. Schmittgen, T. D. & Livak, K. J. Analyzing real-time PCR data by the comparative CT method. Nat. Protoc. 3, 1101–1108 (2008).

65. O’Leary, B. M., Rico, A., McCraw, S., Fones, H. N. & Preston, G. M. The Infiltration-centrifugation Technique for Extraction of Apoplastic Fluid from Plant Leaves Using Phaseolus vulgaris as an Example. JoVE e52113 (2014). doi:doi:10.3791/52113

