## Supplemental Information for "Nanoparticle Cellular Internalization is Not Required for RNA Delivery to Mature Plant Leaves"

[g] Chan-Zuckerberg Biohub, San Francisco, CA 94158, USA.

[§] Authors contributed equally to this work.

*

**Keywords:** Engineered nanomaterials, *in planta*, gold nanoparticles, siRNA delivery

**Correspondence** should be addressed to M.P.L. Mailing address: 492 Stanley Hall, Berkeley, CA 94702, USA.

As a general note, we are working within the confines of a diffraction-limited system when analyzing Cy3 and GFP colocalization under confocal microscopy. Our TEM findings suggest that colocalization values measured in diffraction-limited confocal microscopy represent both cellular internalization and also NP cell wall association, and do not distinguish between the two. While confocal microscopy can inform the accumulation and presence of AuNP in the proximity of plant cells, we are resolution-limited and cannot distinguish if the AuNP are associating with the outer cell wall, between the cell wall and cell membrane, or within the cell. Considering the Abbe diffraction limiting the observation of sub-wavelength features, traditional confocal microscopy alone cannot localize NPs into plant cells[1]. As such, other techniques such as TEM were used to track the passage of AuNP within plant tissue.

**Supplementary Table S1 |** Sequences of oligonucleotides used in this study. Primers used for RT-qPCR were designed using PrimerQuest Tool. The GFP, EF-1, and NbrbohB primer sets produce an amplicon length of 123 bp, 160 bp, and 228 bp respectively.

| Name | Sequence (5'-3') |
|---|---|
| **SH-DNA** | TAC ACG CAT CCT TAG AAA AAA AAA A-SH |
| **Cy3-DNA** | CTA AGG ATG CGT GTA-Cy3 |
| **SH-sense siRNA** | SH-AAA AAA AAA ArGrG rUrGrA rUrGrC rArArC rArUrA rCrGrG rArAT T |
| **Antisense siRNA** | rUrUrC rCrGrU rArUrG rUrUrG rCrArU rCrAC C |

| | |
|---|---|
| **fGFP** | AGT GGA GAG GGT GAA GGT GAT G |
| **rGFP** | GCA TTG AAC ACC ATA AGA GAA AGT AGT G |
| **fEF1** | TGG TGT CCT CAA GCC TGG TAT GGT TG |
| **rEF1** | ACG CTT GAG ATC CTT AAC CGC AAC ATT CTT |
| **SH-sense NT-siRNA** | SH-AAA AAA AAA ArUrA rArGrG rCrUrA rUrGrA rArGrA rGrArU rArCT T |
| **Antisense NT-siRNA** | rGrUrA rUrCrU rCrUrU rCrArU rArGrC rCrUrU rATT |
| **fNbrbohB** | TTT CTC TGA GGT TTG CCA GCC ACC ACC TAA |
| **rNbrbohB** | GCC TTC ATG TTG TTG ACA ATG TCT TTA ACA |

**Supplementary Table S2 |** Assembly of AuNP via the pH-assisted method.

| **AuNP type** | **Concentration** | **Molar Ratio (AuNP:DNA)** | **Centrifugation speed (x 1000g)** |
|---|---|---|---|
| 5nm AuNS | 90 nM | 1:167 | 23 |
| 10nm AuNS | 20 nM | 1:300 | 17 |
| 15nm AuNS | 8 nM | 1:500 | 14 |
| 20nm AuNS | 2.4 nM | 1:800 | 8 |
| AuNR | 2 nM | 1:7500 | 8.5 |

**Supplementary Table S3 |** DLS data of AuNS and NA-AuNS

| **AuNP type** | **Citrate-AuNP** | **DNA-AuNP** | **siRNA-AuNP** |
|---|---|---|---|
| 5 nm AuNS | 11.2±0.4 nm | 15.4±0.7 nm | 17.1±0.7 nm |
| 10 nm AuNS | 14.8±0.1 nm | 20.2±0.6 nm | 19.3±0.3 nm |
| 15 nm AuNS | 18.8±0.2 nm | 27.8±0.5 nm | 25.1±0.4 nm |
| 20 nm AuNS | 23.1±0.4 nm | 30.3±0.3 nm | 32.5±0.4 nm |

**Supplementary Table S4 |** Summary of TEM-quantified AuNR orientation relative to the cell wall

| **Angle of Orientation (°)** | **Number of AuNRs** | **Percentage of AuNR (%)** |
|---|---|---|
| 0 – 10 | 73 | 22.5 |
| 10 – 20 | 36 | 11.1 |
| 20 – 30 | 26 | 8.0 |
| 30 – 40 | 27 | 8.3 |
| 40 – 50 | 25 | 7.7 |
| 50 – 60 | 19 | 5.9 |
| 60 – 70 | 30 | 9.3 |
| 70 – 80 | 42 | 13.0 |
| 80 – 90 | 46 | 14.2 |

**Supplementary Table S5 |** Quantity of DNA or siRNA functionalized on AuNP

| AuNP type | # DNA/AuNP | # siRNA/AuNP |
|---|---|---|
| 5 nm AuNS | 10±1 | 10±2 |
| 10 nm AuNS | 26±2 | 26±10 |
| 15 nm AuNS | 70±6 | 59±18 |
| 20 nm AuNS | 97±4 | 92±14 |
| AuNR | 128±21 | 125±36 |

**Supplementary Table S6 |** Statistical Significances for Nuclease Protection

| Time Elapsed (min) | | Free siRNA | 10 nm AuNS | AuNR |
|---|---|---|---|---|
| | **Column Names** | A | B | C |
| 0 | **% Intact** | 100 | 100 | 100 |
| | **Column comparisons** | | | |
| 10 | **% Intact** | 6.8 | 85.4 | 65.9 |
| | **Column comparisons** | bC | C | |
| 30 | **% Intact** | 6.7 | 75.2 | 62.8 |
| | **Column comparisons** | bc | | |
| 60 | **% Intact** | 3.9 | 67.8 | 62.8 |
| | **Column comparisons** | Bc | | |
| 120 | **% Intact** | 1.8 | 56.4 | 66.2 |
| | **Column comparisons** | BC | | |
| 240 | **% Intact** | 1.0 | 52.1 | 66.9 |
| | **Column comparisons** | Bc | C | |

Means with the same letter have statistically significant differences. Upper-case letters indicate results with $p < 0.05$, and lower-case letters indicate results with $p < 0.001$.

**Fig. S1 | UV-Vis spectra of pristine AuNP and DNA-AuNP confirm DNA-AuNP colloidal stability.**

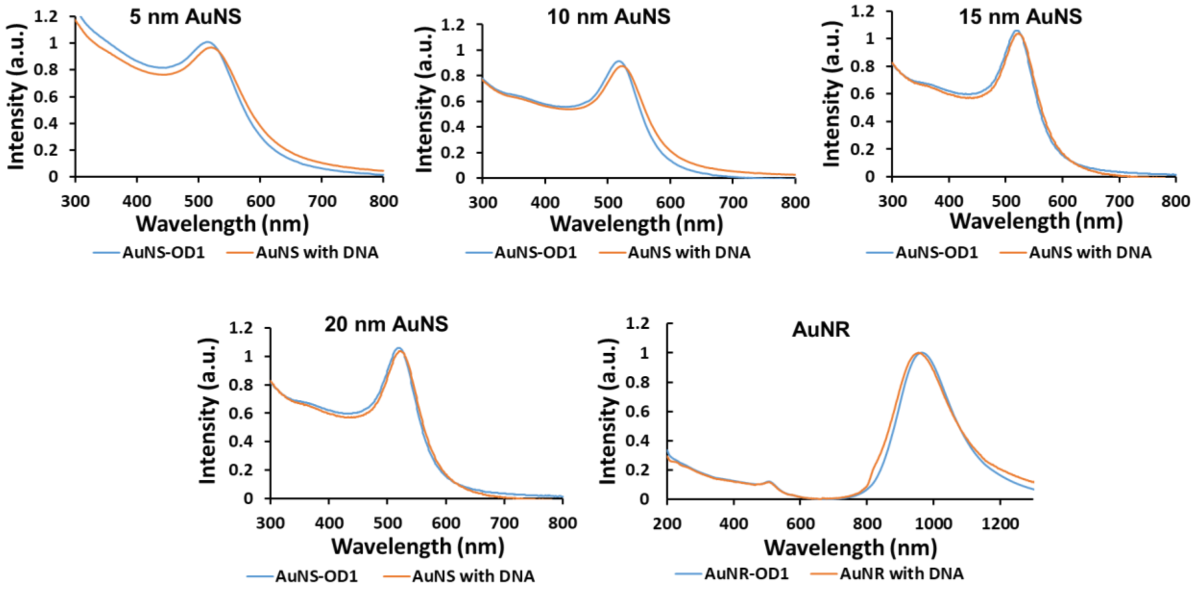

**Fig. S2 | DLS spectra of citrate stabilized AuNS.**

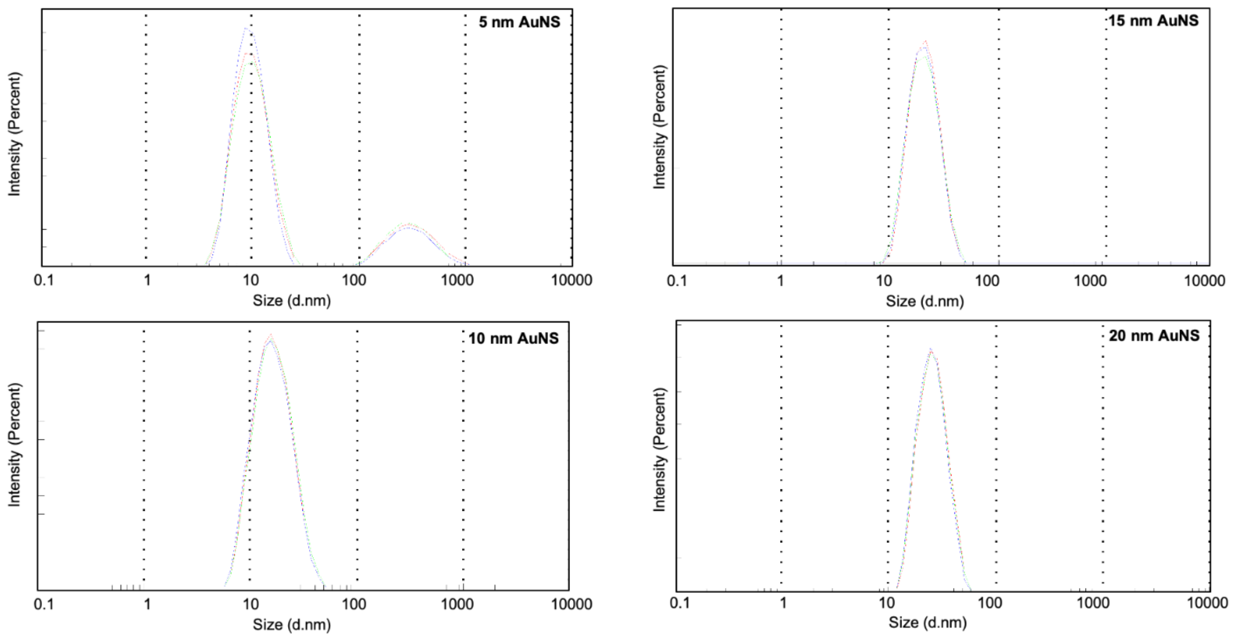

### Fig. S3 | DLS spectra of DNA-AuNS.

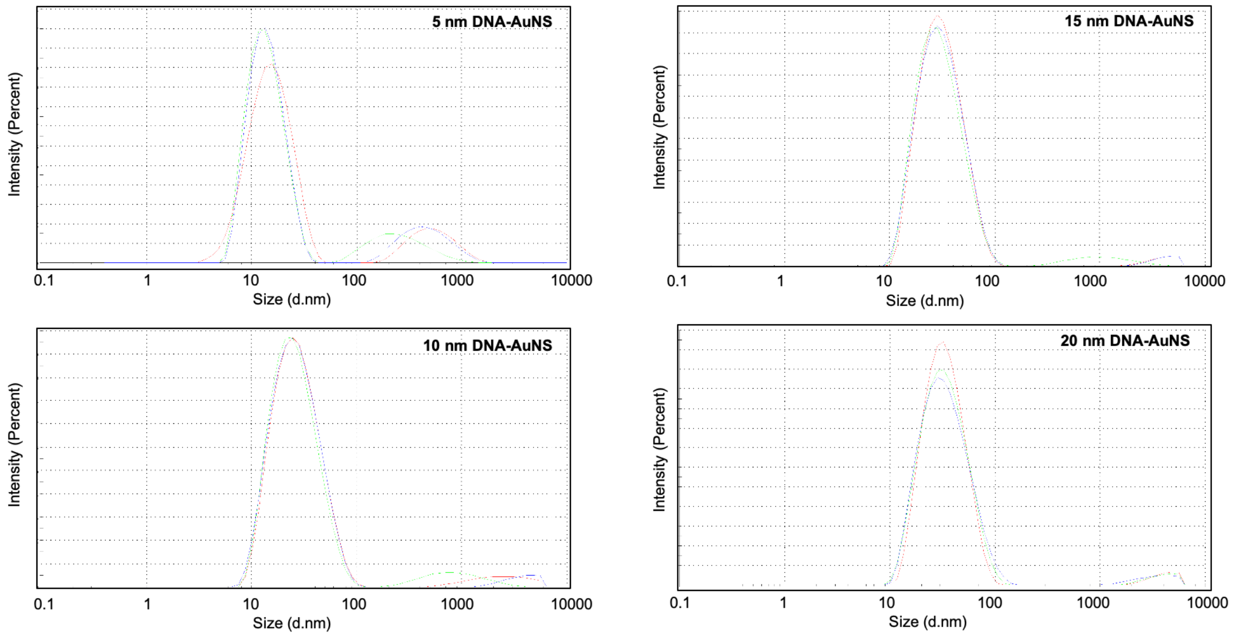

**Fig. S4 | TEM images of DNA-AuNP. Scale Bar: 20 nm for 5 nm AuNS, and 50 nm for other AuNP.**

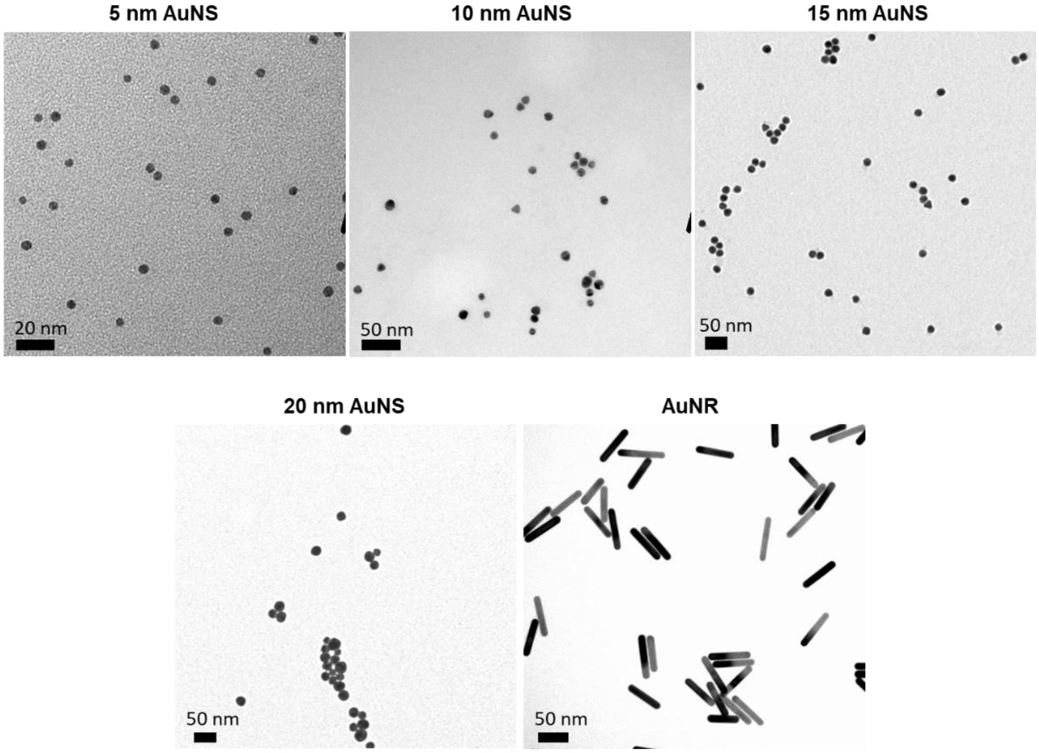

**Fig. S5 | Representative confocal images of 5 nm Cy3-DNA-AuNS infiltrated into *Nicotiana benthamiana* (*Nb*) leaves for various incubation times. Scale bar: 100 μm.**

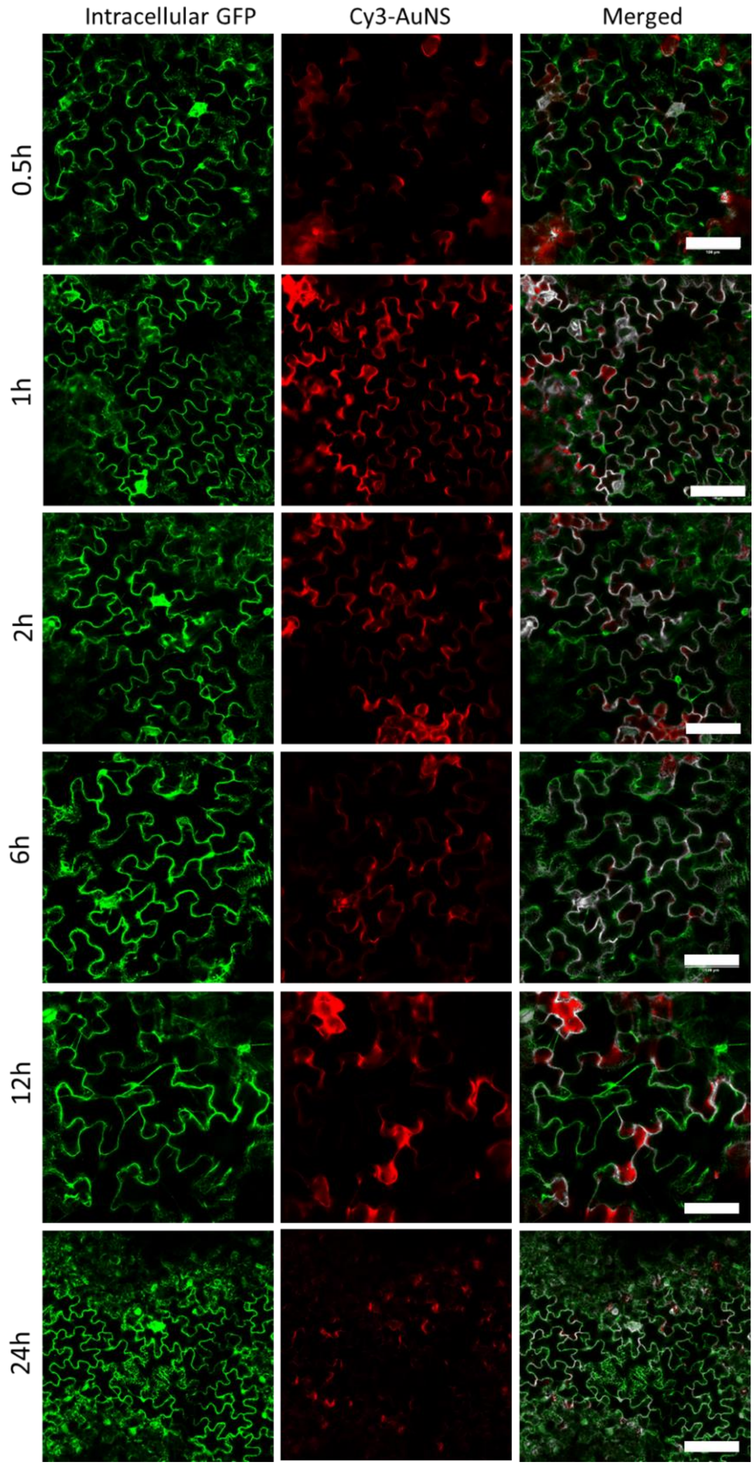

**Fig. S6 | Representative confocal images of 10 nm Cy3-DNA-AuNS infiltrated into *Nb* leaves for various incubation times. Scale bar: 100 μm.**

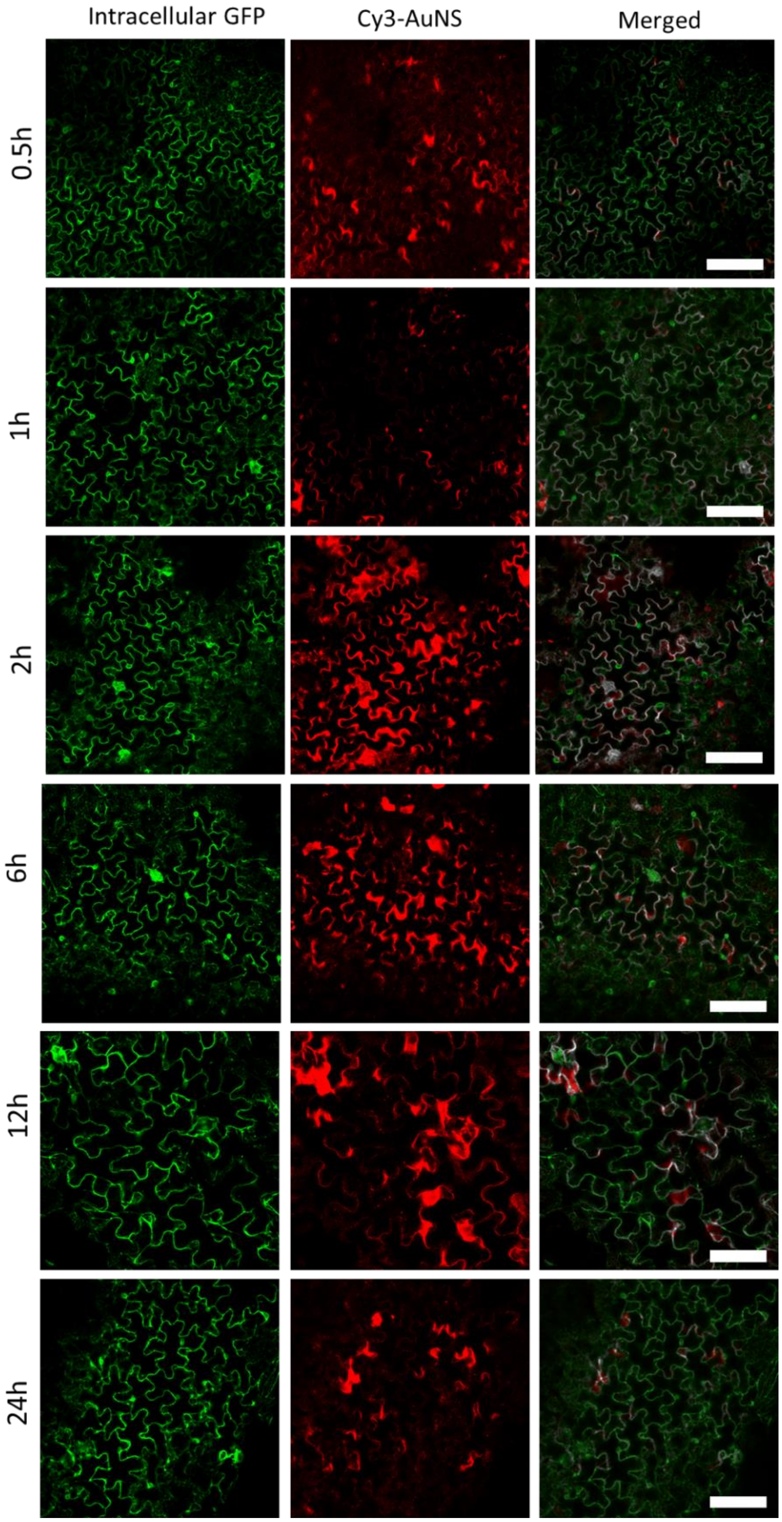

**Fig. S7 | Representative confocal images of 15 nm Cy3-DNA-AuNS infiltrated into *Nb* leaves for various incubation times. Scale bar: 100 μm.**

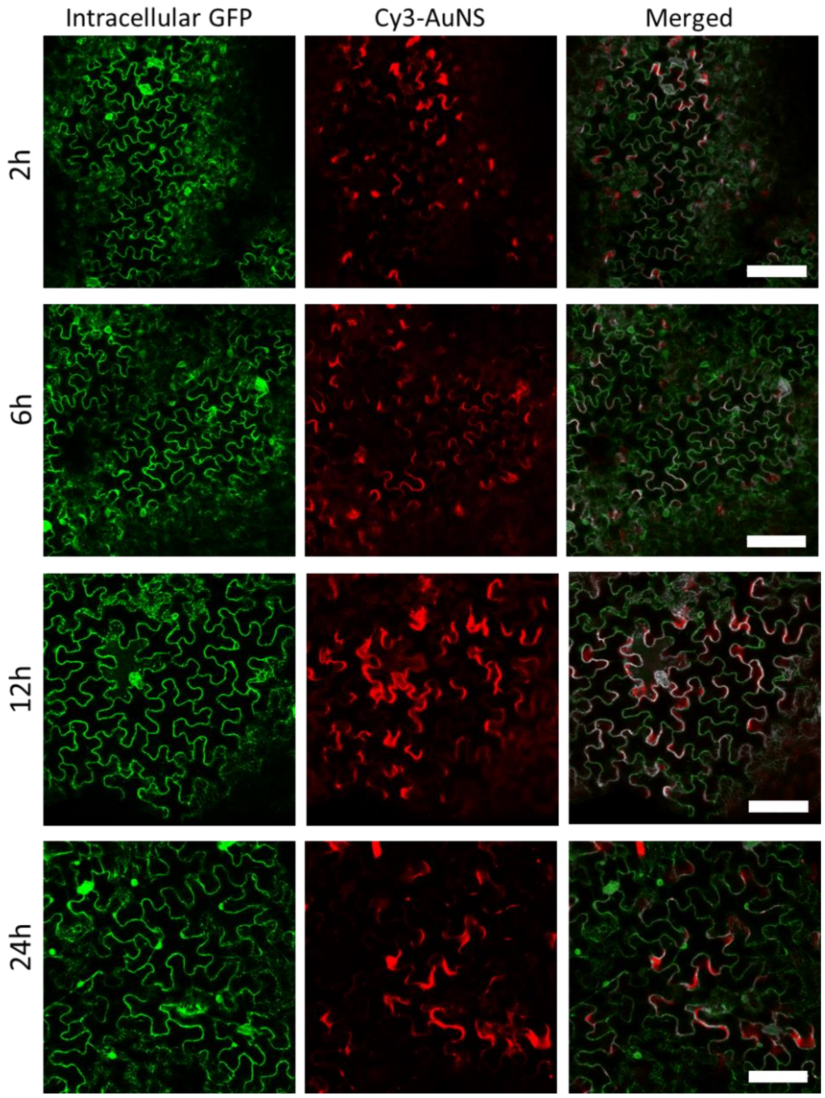

**Fig. S8 | Representative confocal images of 20 nm Cy3-DNA-AuNS infiltrated into *Nb* leaves for various incubation times. Scale bar: 100 µm.**

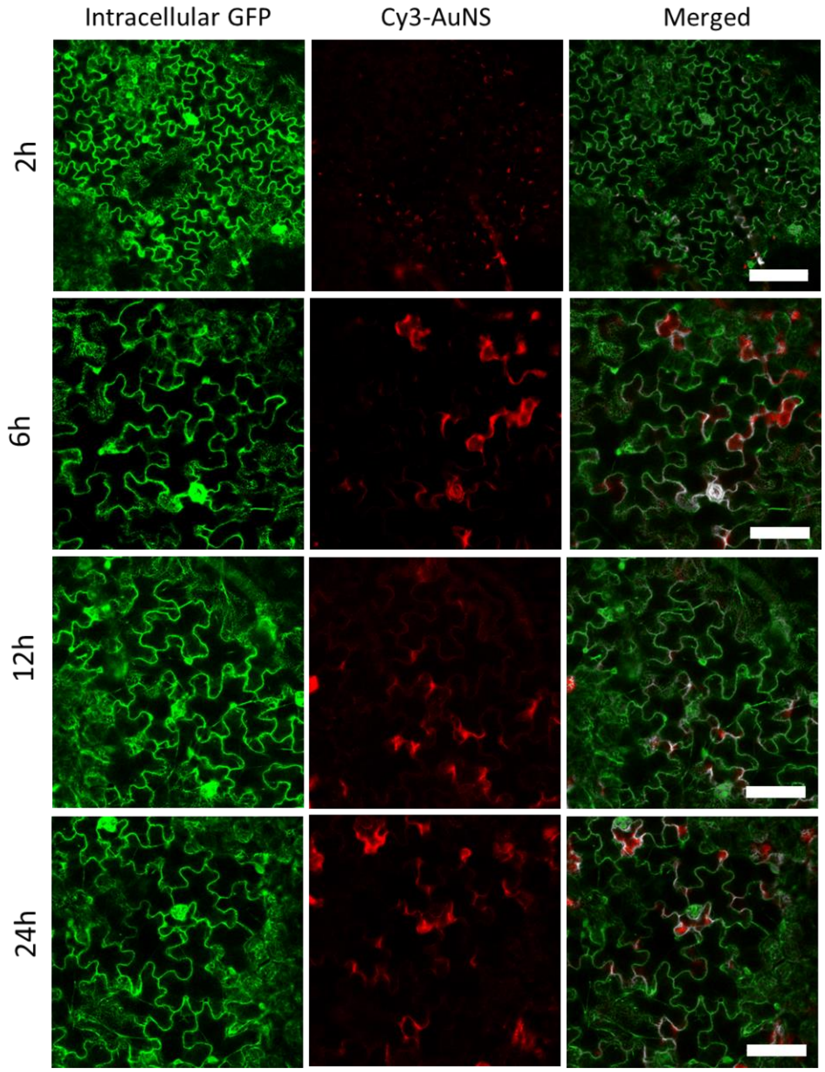

**Fig. S9 | Representative confocal images of Cy3-DNA-AuNR infiltrated into *Nb* leaves for various incubation times. Scale bar: 100 μm.**

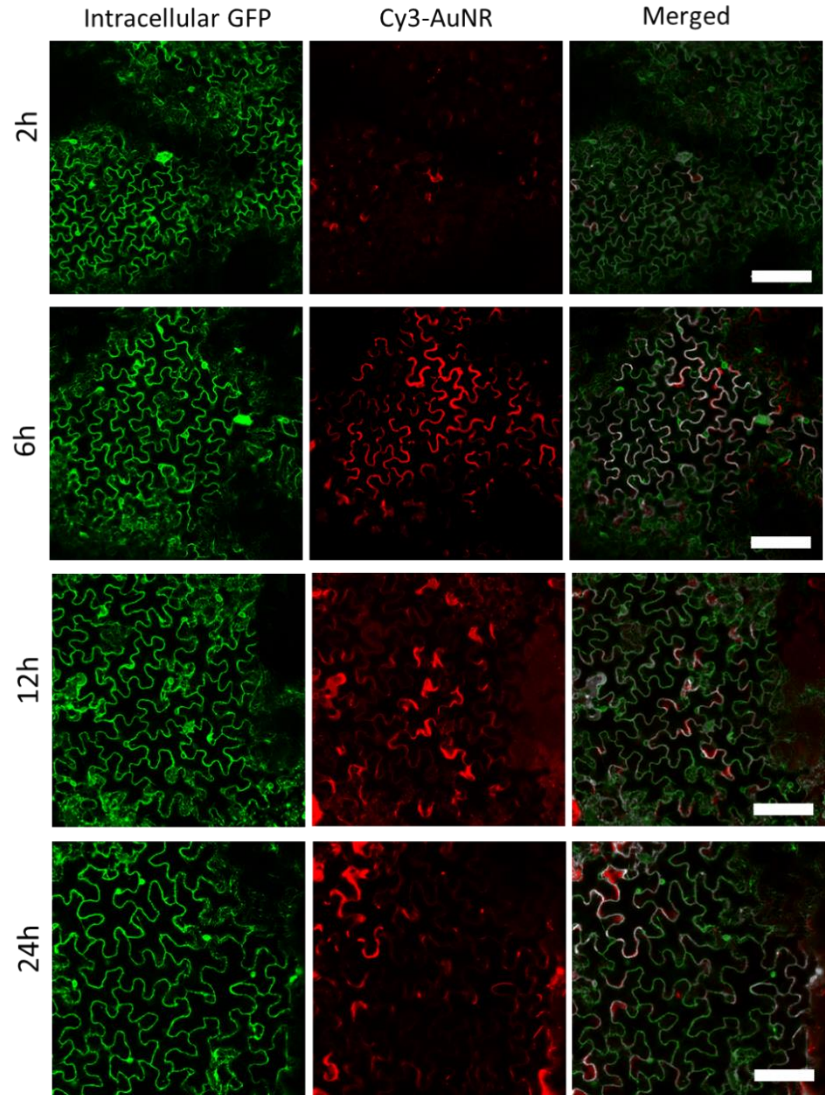

**Fig. S10 | Additional TEM images of DNA-AuNS treated *Nb* leaves. (a) 5 nm DNA-AuNS, (b) 10 nm DNA-AuNS, (c) 15 nm DNA-AuNS, and (d) 20 nm DNA-AuNS.**

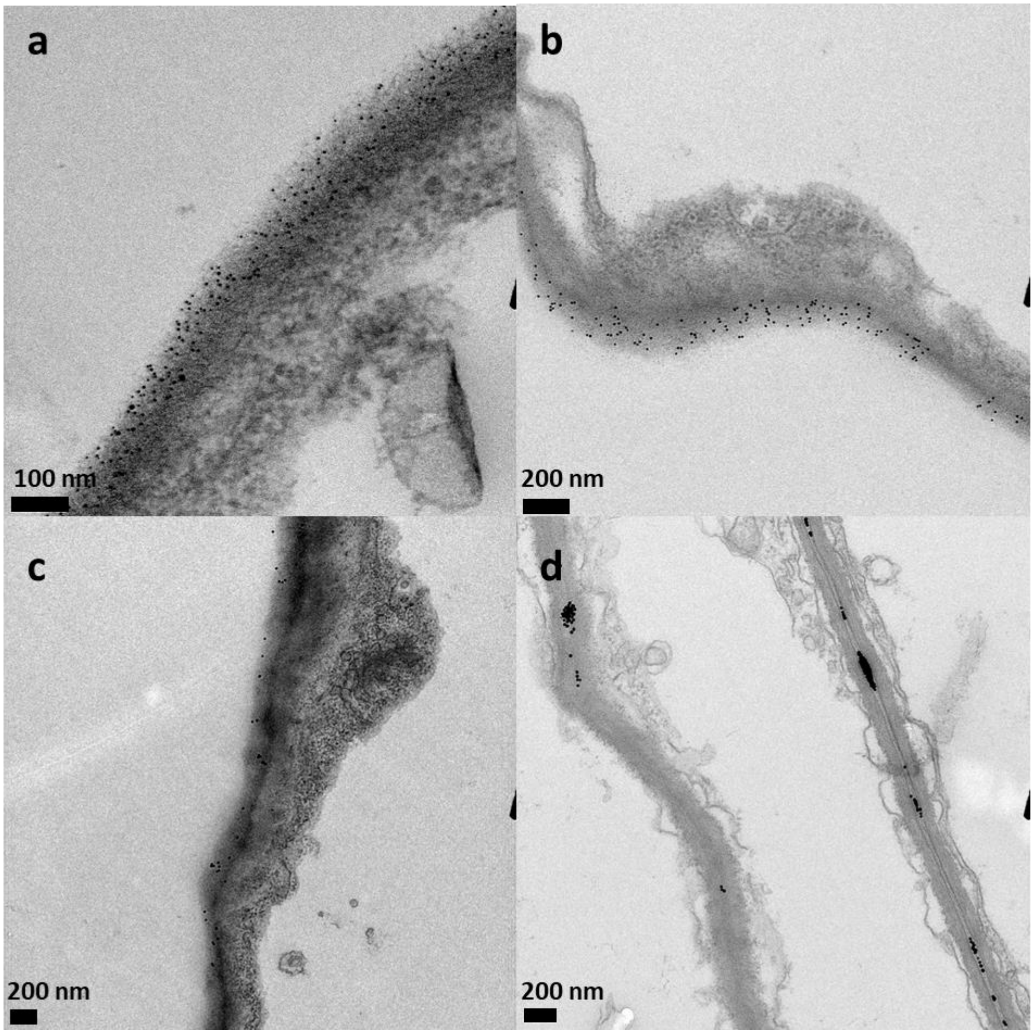

**Fig. S11 | Additional TEM images of DNA-AuNR treated *Nb* leaves.**

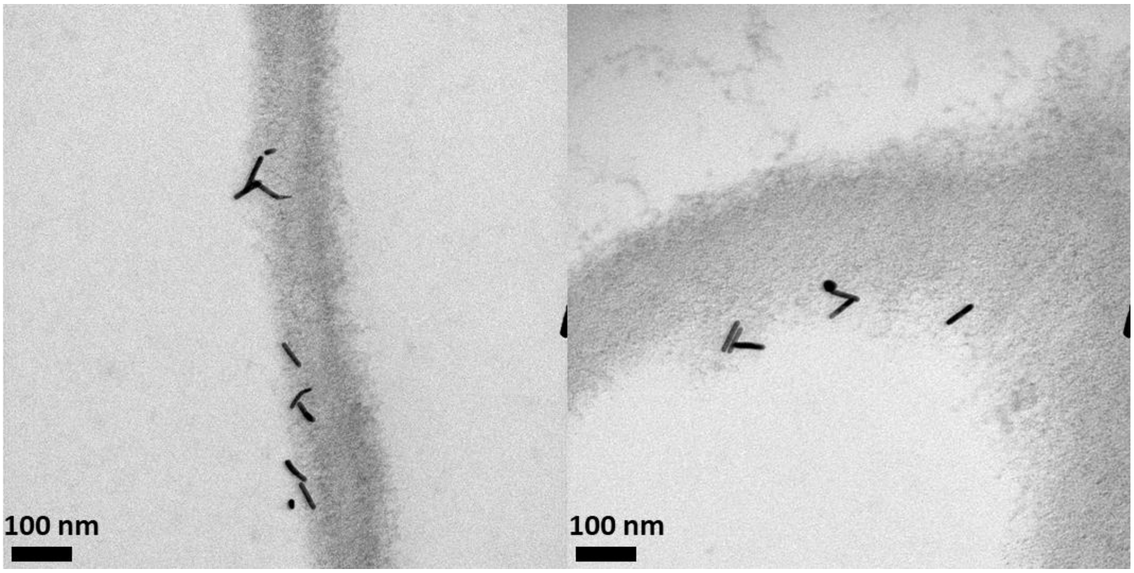

**Fig. S12 | Histogram of AuNR orientations quantified with respect to cell wall tangent. AuNR at 90° are perpendicular to the local region of the cell wall, and AuNR at 0° are parallel to the cell wall. N = 324.**

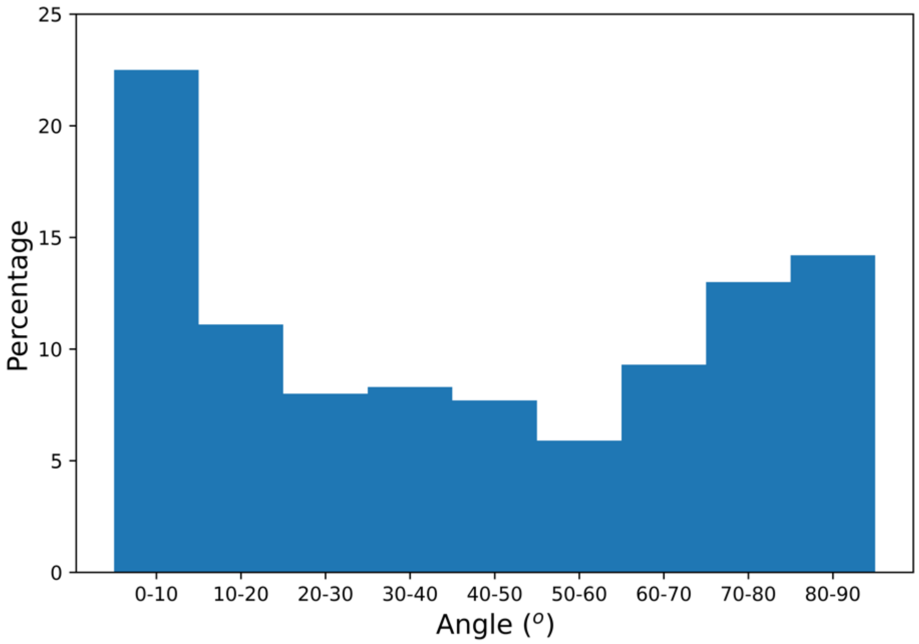

**Fig. S13 | Representative confocal images of Cy3-DNA-AuNP in leaves treated with plant endocytosis inhibitors Ikarugamycin or Wortmanin. Scale bar: 100 μm**

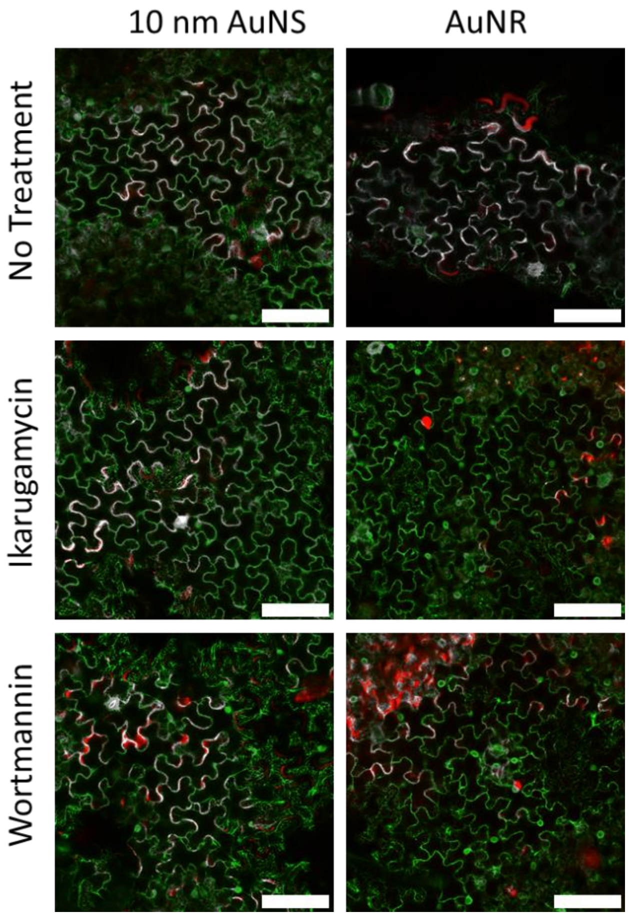

**Fig. S14 | DLS spectra of siRNA-AuNS.**

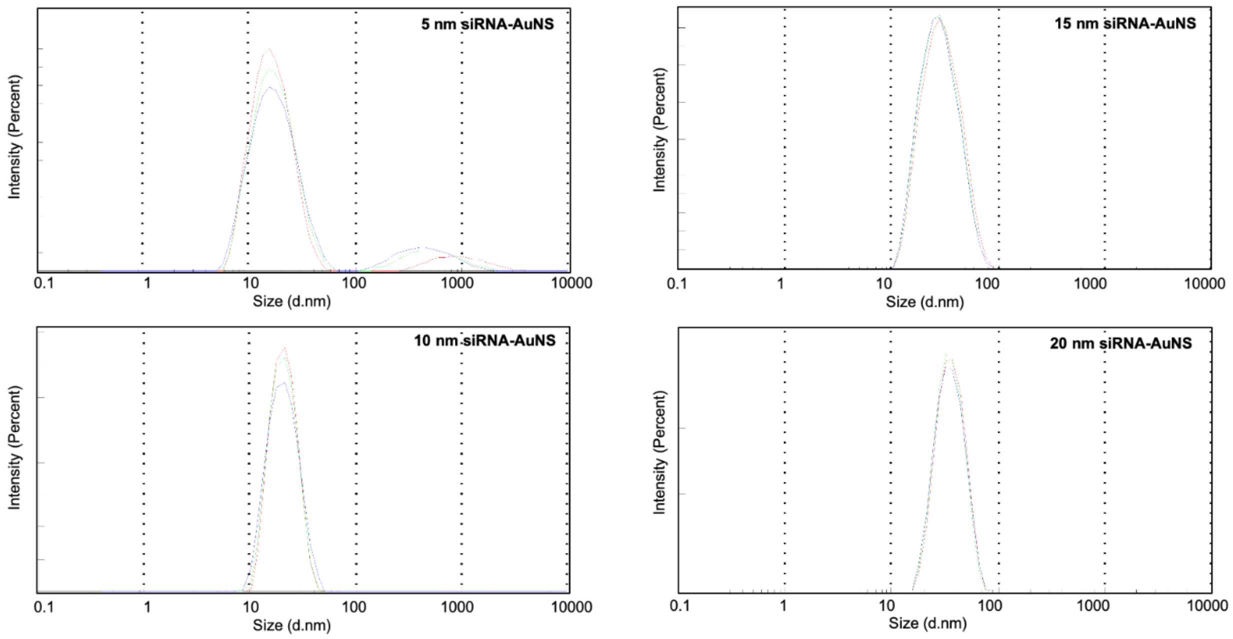

**Fig. S15 | qPCR analysis of GFP gene 1-day post infiltration of non-target siRNA-AuNP.** Buffer, free non-target siRNA (NT-siRNA), or 10 nm NT-siRNA-AuNS were infiltrated into 1-month-old *Nb* leaves and qPCR analysis performed 1-day post-infiltration. There is some variability in GFP mRNA expression levels in both free NT-siRNA and 10 nm NT-siRNA-AuNS samples, though there is no statistically significant difference between the three sample groups. The sequence of NT-siRNA[2] is included in Table S1.

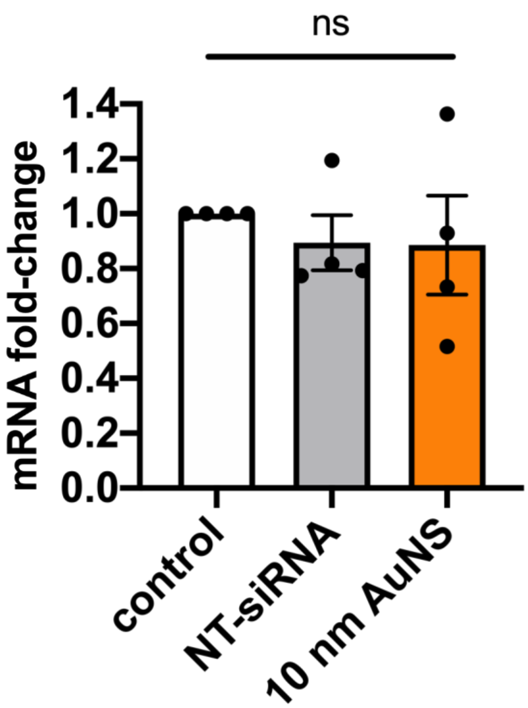

**Fig. S16 | Incubation with plant biofluids induces siRNA desorption from the surface of siRNA-AuNP.** siRNA-AuNP were incubated with water, apoplastic fluid, or lysate on the benchtop. 24 hours post-incubation, the solutions were centrifuged, and supernatants collected to run on a 3% agarose gel. We note the emergence of a band in both the siRNA-AuNP samples incubated with apoplastic fluid and cell lysate that corresponds to the size of free siRNA. This suggests that there is desorption of intact siRNA from siRNA-AuNP in plant biofluids.

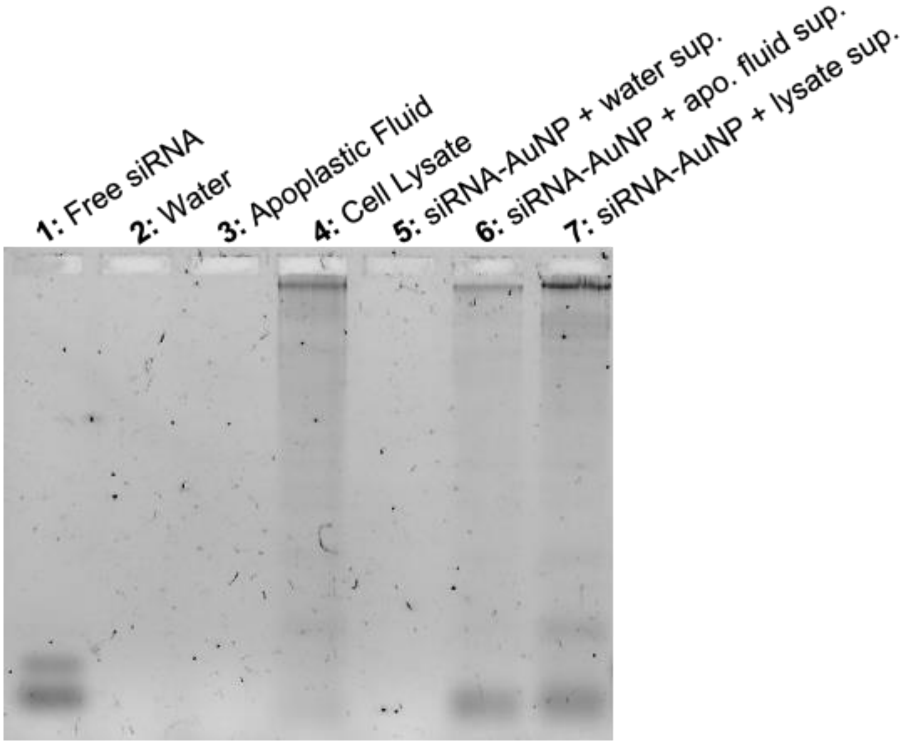

**Fig. S17 | qPCR analysis of *NbrbohB* gene 1-day post-infiltration of siRNA-AuNP.** *NbrbohB* is a *Nb* stress gene that is upregulated in response to various types of stress, including biotic, heat, and mechanical stress[3], and has been used as a proxy for gauging toxicity responses in the presence of nanomaterials [2,4].

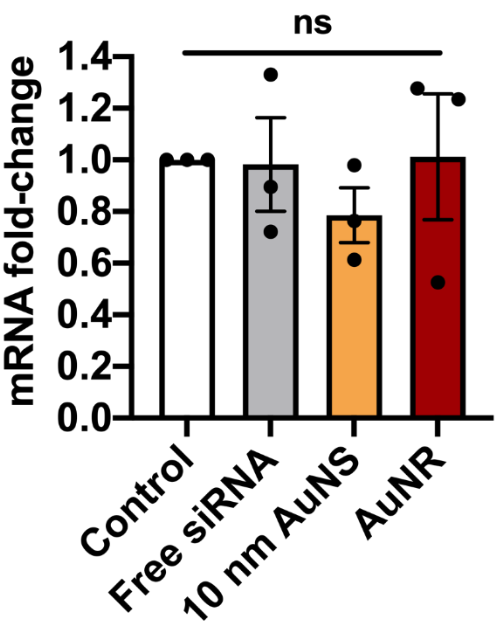

**Fig. S18 | AuNP protect siRNA from endoribonuclease RNase A degradation.** 3% agarose gel of siRNA-loaded on 15nm AuNS or AuNR incubated with RNase A for 1, 6, 12, 18, and 24h, with gel band intensities normalized to siRNA standards and used to calculate extent of siRNA degradation. A two-way ANOVA analysis and Tukey's multiple comparisons test was run. * demonstrates $p<0.05$ when comparing AuNR and free siRNA at a certain timepoint, and + demonstrates $p<0.05$ when comparing AuNR and 15 nm AuNP at a certain timepoint. Error bars represent standard error (n = 3).

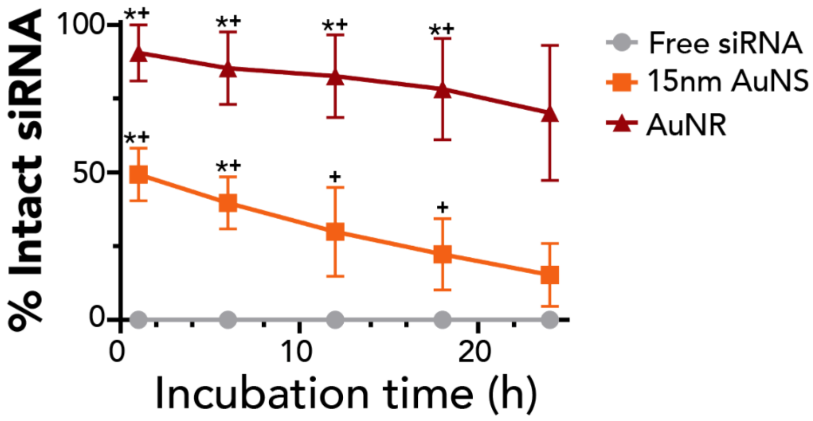

**References**

1. Kuang, C. *et al.* Breaking the diffraction barrier using fluorescence emission difference microscopy. *Sci. Rep.* **3**, 1–6 (2013).

2. Demirer, G. S. *et al.* Carbon nanocarriers deliver siRNA to intact plant cells for efficient gene knockdown. *Sci. Adv.* **6**, eaaz0495 (2020).

3. Yoshioka, H. *et al.* Nicotiana benthamiana gp91phox homologs Nbrboha and Nbrbohb participate in H2O2 accumulation and resistance to Phytophthora infestans. *Plant Cell* **15**, 706–718 (2003).

4. Demirer, G. S. *et al.* High aspect ratio nanomaterials enable delivery of functional genetic material without DNA integration in mature plants. *Nat. Nanotechnol.* **14**, 456–464 (2019).
